# Loss of the Familial Dysautonomia gene *Elp1* in cerebellar granule cell progenitors leads to ataxia in mice

**DOI:** 10.1101/2024.03.27.586801

**Authors:** Frederik Manz, Patricia Benites Goncalves da Silva, Mackenna E. Schouw, Chiara Lukasch, Luca Bianchini, Laura Sieber, Jesus Garcia-Lopez, Shiekh Tanveer Ahmad, Yiran Li, Hong Lin, Piyush Joshi, Lisa Spänig, Magdalena Radoš, Mykola Roiuk, Mari Sepp, Marc Zuckermann, Paul A. Northcott, Annarita Patrizi, Lena M. Kutscher

## Abstract

Familial Dysautonomia (FD) is an autosomal recessive disorder caused by a splice site mutation in the gene ELP1, which disproportionally affects neurons. While classically characterized by deficits in sensory and autonomic neurons, neuronal defects in the central nervous system have been described. ELP1 is highly expressed in the normal developing and adult cerebellum, but its role in cerebellum development is unknown. To investigate the cerebellar function of Elp1, we knocked out Elp1 in cerebellar granule cell progenitors (GCPs) and examined the outcome on animal behavior and cellular composition. We found that GCP-specific conditional knockout of Elp1 (Elp1^cKO^) resulted in ataxia by 8 weeks of age. Cellular characterization showed that the animals had smaller cerebella with fewer granule cells. This defect was already apparent 7 days after birth, when Elp1^cKO^ animals also exhibited fewer mitotic GCPs and shorter Purkinje dendrites. Through molecular characterization, we found that loss of Elp1 was associated with an increase in apoptotic cell death and cell stress pathways in GCPs. Our study demonstrates the importance of ELP1 within the developing cerebellum, and suggests that Elp1 loss in the GC lineage may also play a role in the progressive ataxia phenotypes of FD patients.

## INTRODUCTION

Familial Dysautonomia (FD), also called Riley-Day syndrome or hereditary sensory and autonomic neuropathy type III (HSAN III), is an autosomal-recessive degenerative disorder of the sensory and autonomic nervous system (Rotthier et al. 2012). Neuronal deficits have also been reported in the central nervous system (Mendoza-Santiesteban et al. 2012; Gutiérrez et al. 2015). Hallmark symptoms of FD include temperature and pain insensitivity, inability to cry, cardiovascular instability, retinal degeneration and progressive ataxia (Rotthier et al. 2012; González-Duarte et al. 2023); only 50% of patients reach age 40 (Axelrod et al. 2002). Genetically, FD is caused by homozygous splice-site mutation in the *ELP1* (Anderson et al. 2001), which results in tissue-specific skipping of exon 20 and selective loss of ELP1 protein (Slaugenhaupt et al. 2001), especially in neurons (Cuajungco et al. 2003). *ELP1* encodes the scaffolding subunit of the highly conserved six-subunit Elongator complex, which modifies specific tRNAs with 5-carboxymethyl derivatives (xcm^5^), ensuring anticodon-codon pairing and effective protein translation (Karlsborn et al. 2014; Nedialkova and Leidel 2015).

The cerebellum primarily coordinates motor function and balance, and dysfunction of the cerebellum is often associated with progressive ataxias (Schmahmann 2019). In FD patients, *ELP1* is preferentially mis-spliced in the cerebellum compared with other organs (Hims et al. 2007), suggesting that ELP1 protein levels may also be decreased in the cerebellum of FD patients. Among healthy tissues, *ELP1* gene expression is the highest in the cerebellum compared to other brain regions and organs in humans (Anderson et al. 2001; Waszak et al. 2020). However, the role of *ELP1* in the cerebellum and how loss of *ELP1* may contribute to ataxia is unexplored.

To investigate the function of *Elp1* in the cerebellum and determine whether loss of *Elp1* may contribute to ataxia, we used an *Atoh1-Cre* driven conditional knockout approach to eliminate *Elp1* in developing granule cell progenitor (GCP) cells (*Elp1*^*cKO*^). *Elp1*^*cKO*^ animals developed ataxia by 8 weeks of age. While the overall cytoarchitecture of *Elp1*^*cKO*^ cerebellum was unaffected, *Elp1*^*cKO*^ animals had fewer GCs at 4 and 12 weeks of age, resulting in smaller cerebella compared with littermate controls. At postnatal day 7 (P7), *Elp1*^*cKO*^ animals had fewer mature GCs, decreased expression of the glutamatergic transporter VGluT1, and shorter Purkinje dendrites. These cellular phenotypes were likely the direct result of increased apoptosis at P7. Our study suggests that *Elp1* loss in the GC lineage may contribute to the ataxic symptoms seen in FD patients.

## RESULTS

### ELP1 is highly expressed in proliferating granule cell progenitors and protein levels are downregulated upon differentiation

Previous studies have shown that *ELP1* exhibits the highest expression in the postnatal cerebellum throughout the development of major organs in humans, and in mice its expression gradually increases postnatally in the GC lineage cells (Waszak et al. 2020). To further understand *ELP1* expression patterns during human and mouse cerebellum development, we examined *ELP1* levels across developmental stages and cell types. Bulk RNA-sequencing data (Cardoso-Moreira et al. 2019) indicates a trend of increasing *ELP1* expression in the cerebellum postnatally in humans, whereas a decline is observed in the mouse (**Supplemental Fig. S1A,B**); however, the murine cerebellum still maintains highest *Elp1* expression levels compared to other organs in adult mice. To compare gene expression between species at the single-cell level, we examined cell-type-specific expression using singlenucleus RNA-sequencing atlases of the developing human and mouse cerebellum (Sepp et al. 2024). *ELP1*/ *Elp1* is broadly expressed across most cell types during cerebellum development (**Supplemental Fig. S1C,D**). Similar to bulk expression in the mouse, overall *Elp1* expression at the single-cell level decreased upon birth in the majority of cell types, but expression was still sustained to adulthood in many cell types, including the GC, interneuron, and glial lineages (**Fig. 1A**; **Supplemental Fig. S1C**). In humans, *ELP1* expression was maintained during development or increased towards adulthood in the GC, interneuron and glial lineages (**Fig. 1B**; **Supplemental Fig. S1D**). Overall, despite varied developmental dynamics across species, *ELP1*/*Elp1* is broadly expressed in the mouse and human cerebellum and in granule cells its expression is sustained from early embryonic stages to adulthood.

**Figure 1.**
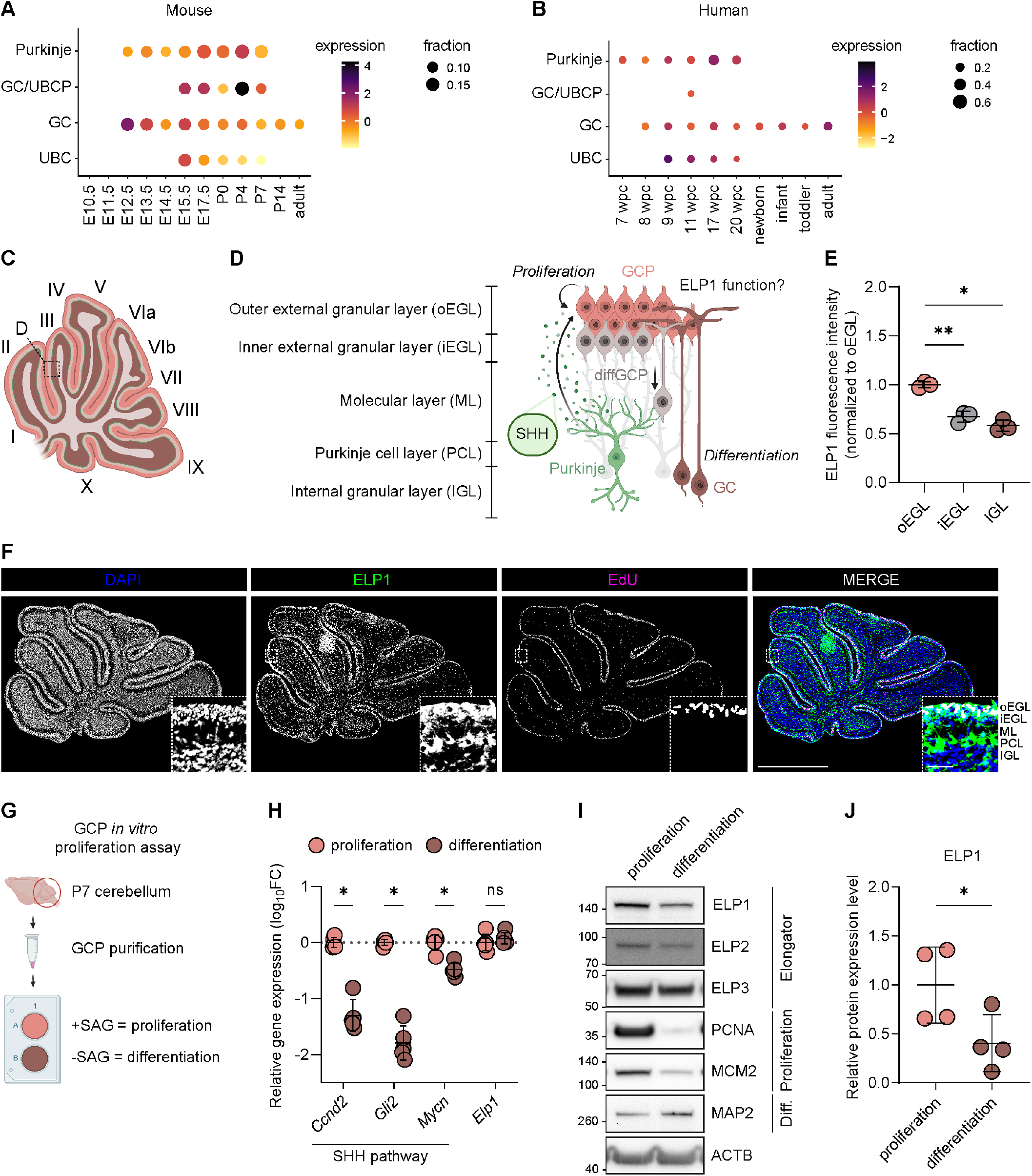
ELP1 is expressed in proliferating granule cell progenitors during cerebellar development. (**A-B**) Expression of *Elp1/ELP1* in Purkinje and granule cell lineage cells across different time points of murine (**A**) and human (**B**) cerebellar development. Dot size and color indicate fraction of cells and mean expression of *Elp1*/*ELP1* averaged across biological replicates, respectively. Extended dot plot in Supplemental Fig. S1C,D. GC/UBCP, granule cell/unipolar brush cell progenitor. (**C**) Cartoon showing sagittal cut of P7 murine cerebellum. (**D**) Cartoon showing layered cytoarchitecture and stages of GC lineage development in P7 mouse cerebellum. (**E**) Median ELP1 fluorescence intensity in outer external granular layer (oEGL), inner external granular layer (iEGL) and internal granular layer (IGL) across lobes III, VI, IX and X. Paired *t*-tests. Mean ± SD. *N* = 3 mice, *n =* 4 lobes. (**F**) Representative immunofluorescence image of DAPI (blue), ELP1 (green) and EdU (magenta) in P7 *Elp1*^WT^ cerebellum. Scale bar, 1 mm. Inset, increased magnification of single z-slice of lobe III. Scale bar, 50 μm. ML, molecular layer. PCL, Purkinje cell layer. (**G**) Cartoon showing GCP *in vitro* proliferation assay. SAG, Smoothened agonist. (**H**) Relative mRNA expression (quantitative PCR) of *Ccnd2, Gli2, Mycn* and *Elp1* of GCPs cultured with SAG (orange, proliferation) or without SAG (brown, differentiation), normalized to *Tbp1*. Multiple Mann-Whitney *U* tests. *, *P*<0.005. Mean ± SD. *N* = 5. SHH, Sonic hedgehog. (**I**) Immunoblot validation of ELP1, ELP2, ELP3, PCNA, MCM2, and MAP2 protein in GCPs cultured with SAG (proliferation) or without SAG (differentiation). ACTB was used as loading control. (**J**) Quantification of ELP1 protein abundance in P7 GCP lysates normalized to ACTB. Unpaired *t*-test. Mean ± SD. *N* = 4.

Given the high proportion of GCs in the postnatal cerebellum, high *ELP1* expression levels in GCs, and potential link to ataxia in FD patients, we first investigated protein expression of ELP1 in the GC lineage of the normal developing murine cerebellum (Fig. 1C). In the first two postnatal weeks of mice, Purkinje cells (PCs) secrete SHH and stimulate proliferation of GCPs in the outermost part of the external granular layer (EGL; Fig. 1D; Consalez et al. (2021)). Once GCPs exit cell cycle, they migrate inwards and differentiate into mature GCs, forming the internal granular layer (IGL). At P7, both EGL and IGL are present, which allows us to examine both proliferating GCPs and mature GCs. Therefore, we injected EdU into wild type P7 pups to mark actively cycling cells 2h, and performed immunofluorescence staining against ELP1 and EdU on brains harvested 2h after injection (Fig. 1E,F). We found that proliferating GCPs in the outer EGL expressed ELP1 higher than differentiating GCPs in the inner EGL or mature GCs in the IGL (Fig. 1E,F).

To identify the role of ELP1 in proliferating GCPs, we first validated whether ELP1 expression correlates with cell cycle using an *in vitro* proliferation assay (**Fig. 1G**). Briefly, we isolated GCPs from P7 mice and examined their response to SHH pathway activation by exposure to Smoothened agonist (SAG) *in vitro*. SAG treatment maintains the proliferation of purified GCPs, whereas withdrawal terminates proliferation and induces differentiation after 72h *in vitro* (Wechsler-Reya and Scott (1999); **Supplemental Fig. S2A,B**). As expected, proliferative (SAG-treated) GCPs expressed higher levels of SHH pathway genes *Ccnd2, Gli2, Mycn, Gli1, Ccnd1, Ptch1* and the hallmark GCP transcription factor *Atoh1* compared with differentiating (untreated) GCPs (**Fig. 1H**; **Supplemental Fig. S2A,C**). In contrast, *Elp1* gene expression remained unchanged in proliferating and differentiating GCPs (**Fig. 1H**). At the protein level, however, proliferating GCPs expressed higher levels of ELP1 than differentiated GCs (**Fig. 1I,J**; **Supplemental Fig. S2D,E**). Protein levels of ELP2, another component of the Elongator core complex, were also reduced upon differentiation, but protein levels of ELP3, the catalytic subunit of the Elongator complex, were not reduced (**Fig. 1I**; **Supplemental Fig. S2F,G**). Altogether, ELP1 shows elevated expression in proliferating GCPs, which prompted us to next investigate the consequences of *Elp1* loss in cerebellar GCPs.

### Conditional Elp1 knockout reduces ELP1 and Elongator in GCPs

To investigate the functional relevance of *Elp1* during GC development, we used a genetically engineered mouse strain to generate Cre-inducible excision of exon 4 of *Elp1* in *Atoh1*-positive GCPs (**Fig. 2A**; hereafter called *Elp1*^cKO^). Immunofluorescence staining confirmed a specific reduction of ELP1 in GCPs, but not in PCs in P7 *Elp1*^cKO^ mice (**Fig. 2B**; **Supplemental Fig. S3A**). While ELP1 was absent from the EGL of the anterior and central lobules of *Elp1*^cKO^ cerebella, ELP1 was present in the nodular lobules IX and X, where *Atoh1-Cre* transgene activity is known to be markedly reduced (Pan et al. 2009). We confirmed loss of ELP1 protein in purified *Elp1*^cKO^ GCPs compared with littermate control GCPs (**Fig. 2C,D**). *Elp1*^cKO^ GCPs also showed reduced levels of ELP2 and ELP3 (**Fig. 2C,E,F**), indicating the destabilization and degradation of the core Elongator complex upon ELP1 loss (**Fig. 2G**; Xu et al. (2015)). Despite higher ELP1 expression in proliferating GCPs (**Fig. 1I,J**), proliferation *in vitro* as assessed by EdU incorporation and PCNA protein levels (**Fig. 2H**; **Supplemental Fig. S3B**) was similar in *Elp1*^cKO^ GCPs compared with sibling controls; however, MCM2 protein levels were slightly increased (**Supplemental Fig. S3C**). Response to SHH pathway stimulation was unchanged in *Elp1*^cKO^ GCPs (**Fig. 2I**; **Supplemental Fig. S3D**). The total number of GCPs purified from *Elp1*^cKO^ animals were reduced, however (**Fig. 2J**), suggesting loss of viable GCPs in *Elp1*^cKO^ cerebella. In summary, loss of *Elp1* destabilized the Elongator complex in proliferating GCPs, allowing us to use this mouse model to examine the effect of *Elp1* loss on animal behavior.

**Figure 2.**
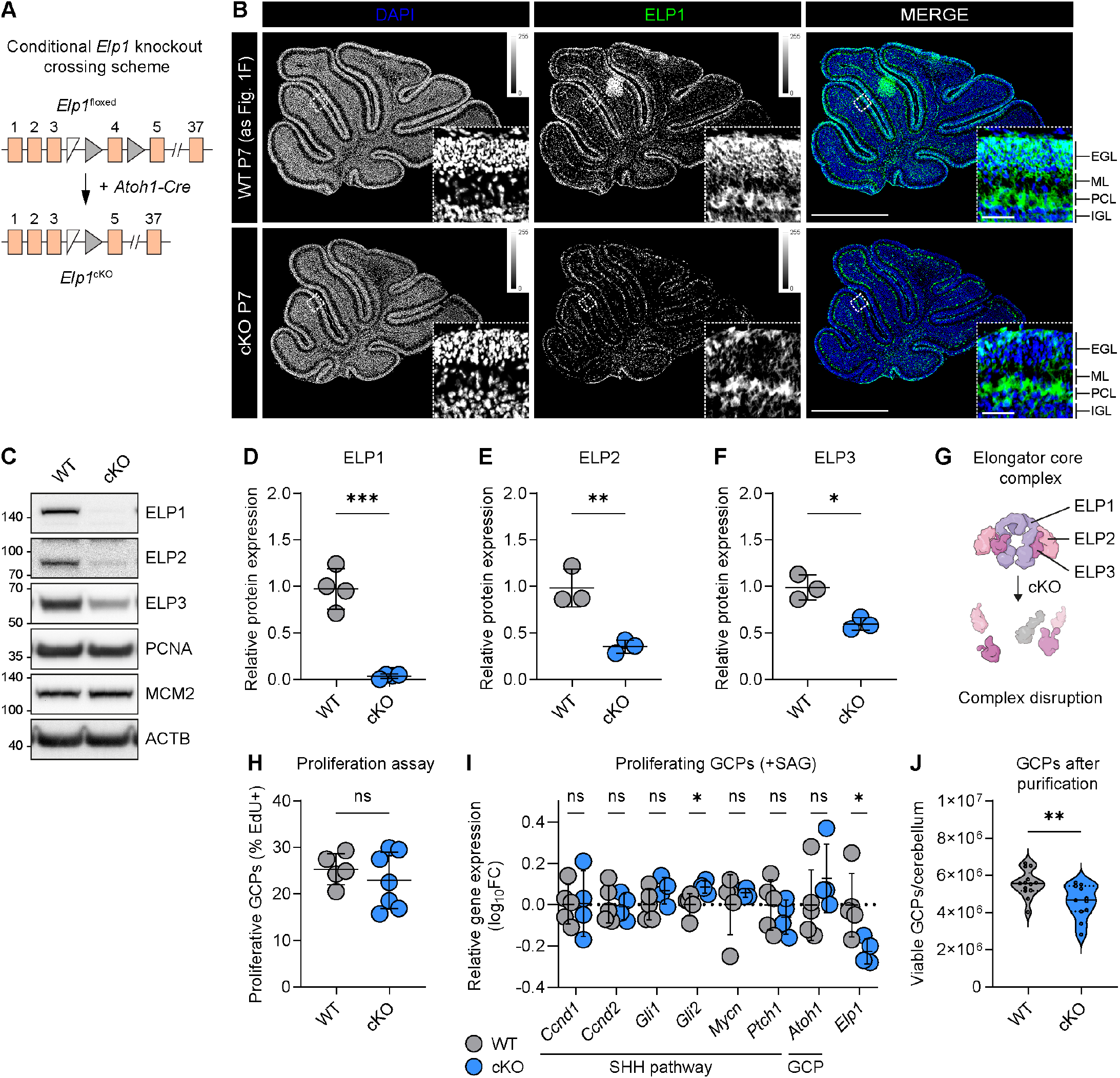
GCP-specific *Elp1* knockout reduces ELP1 and destabilizes the Elongator core complex. (**A**) Schematic showing the crossing scheme of GCP-specific *Elp1* knockout, adapted from Chaverra et al. (2017). (**B**) Representative immunofluorescence imaging of DAPI (blue) and ELP1 (green) expression in P7 cerebella of *Elp1*^WT^ (top row) and *Elp1*^cKO^ (bottom row) littermates. Elp1^WT^ cerebellum is the same as in Fig. 1F. Single z-slice. Scale bar, 1 mm. Inset, magnification of lobe III. Scale bar, 50 μm. EGL, external granular layer. ML, molecular layer. PCL, Purkinje cell layer. IGL, internal granular layer. (**C-F**) Immunoblot validation (**C**) and quantification of ELP1 (**D**), ELP2 (**E**) and ELP3 (**F**) protein abundance from *Elp1*^cKO^ and *Elp1*^WT^ GCP lysates. ACTB was used as loading control. Unpaired *t*-tests. Mean ± SD. *N* = ≥ 3 samples/genotype. (**G**) Sketch shows Elongator core complex and its destabilization upon ELP1 loss. Adapted from Garcia-Lopez et al. (2021). (**H**) Percentage of proliferative GCPs (EdU+) after 72h *in vitro* with Smoothened agonist (SAG). Unpaired *t*-test. Mean ± SD. *N* = ≥ 5 samples/genotype. (**I**) Relative mRNA expression (quantitative PCR) of *Ccnd1, Ccnd2, Gli1, Gli2, Mycn, Ptch1* (all SHH pathway genes), *Atoh1* (GCP marker) and *Elp1* of P7 GCPs after 72h *in vitro* with SAG. Multiple Mann-Whitney *U* tests. Mean ± SD. *N* ≥ 4 samples/genotype. (**J**) Total number of viable GCPs isolated from individual *Elp1*^WT^ or *Elp1*^cKO^ cerebella at P7. Unpaired *t*-test. *N* = 13 (WT) and 11 (cKO).

### Elp1^cKO^ mice develop ataxia

In contrast to the embryonic lethality of systemic *Elp1* knockout (Chen et al. 2009; Dietrich et al. 2011), *Elp1*^cKO^ mice survived to adulthood and developed an insecure, unsteady gait, similar to FD-specific mouse models (Macefield et al. 2011; Macefield et al. 2013; Morini et al. 2023). To assess ataxia onset and progression, we performed a tailored battery of behavioral tests to assess general animal health, muscle strength, balance, motor coordination and gait over time (**Fig. 3A**). Behavioral analysis was performed blinded to genotype and included both sexes. First, we determined the overall health of the *Elp1*^cKO^ mice by a SHIRPA primary screen at week 7 (see Methods for details), followed by movement and strength assessments using a rotarod assay, grip strength assay and gait analysis at weeks 8, 10 and 12 (**Supplemental Tables S2-S4**). In the SHIRPA screen at week 7, male *Elp1*^cKO^ mice had similar scores as their *Elp1*^WT^ littermates while female *Elp1*^cKO^ mice had slightly worse scores than their *Elp1*^WT^ littermates (**Fig. 3B**). Body weight and forelimb grip strength remained mostly unchanged during the experimental period, although male mice showed a slight decrease at week 12 (**Fig. 3C,D**). Motor performance of *Elp1*^cKO^ mice was significantly decreased across both sexes and all time points (**Fig. 3E**), indicating a compromised motor performance compared to their control littermates.

**Figure 3.**
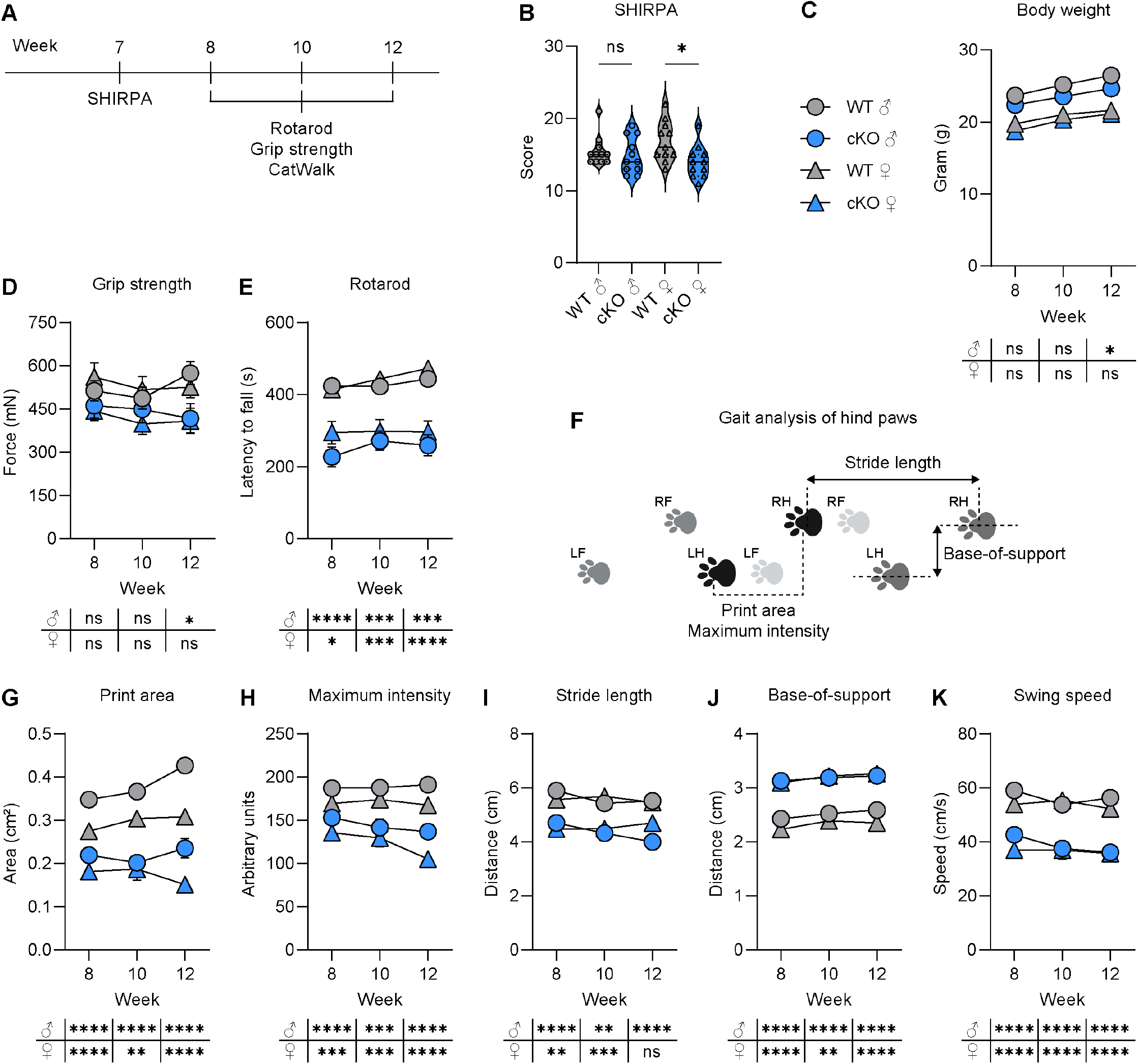
Adult *Elp1*^cKO^ mice have impaired motor coordination and gait ataxia. (**A**) Schematic overview of the motor behavior tests. (**B**) SHIRPA score of male (circles) and female (triangles) *Elp1*^WT^ (gray) and *Elp1*^cKO^ (blue) animals at week 7. (**C**) Average body weight of animals across sex and time points. (**D**) Average grip strength across sex and time points. (**E**) Average latency to fall in the rotarod test across sex and time points. (**F**) Graphical illustration of selected hind paw gait parameters. LF, left front paw. RF, right front paw. LH, left hind paw. RH, right hind paw. Adapted from Pitzer et al. (2021). (**G-K**) Quantification of hind paw gait parameters across sex and time points: print area (**G**), maximum intensity (**H**), stride length (**I**), base-of-support (**J**) and swing speed (**K**). (**B-E; G-K**) Mann-Whitney *U* tests. Mean ± SEM. *N* = 11 mice per time point, genotype and sex. Raw data is available in Supplemental Tables S2-S4.

To characterize the onset and progression of the ataxic gait, we performed a gait analysis using the CatWalk XT system (**Fig. 3F**; Heinzel et al. (2020); Pitzer et al. (2021)). Multiple gait parameters were disrupted in *Elp1*^cKO^ mice compared with *Elp1*^WT^ mice littermates, in particular in the hind paws (**Fig. 3F-K**). We detected a reduction of the area and the maximum intensity of the paw print, which indicated that *Elp1*^cKO^ animals placed less weight on a reduced area of their hind paws (**Fig. 3G,H**). These parameters were decreased in both sexes across all time points.

Stride length and base-of-support are important distance parameters to assess animal gait (Hamers et al. 2006) and are often compromised in patients with cerebellar ataxia (Palliyath et al. 1998; Stolze et al. 2002). *Elp1*^cKO^ mice had a reduced stride length, indicating the distance between consecutive placements of the same paw was decreased (**Fig. 3I**). *Elp1*^cKO^ mice also had an increased base-of-support, such that animals increased the distance between both hind paws to better stabilize themselves (**Fig. 3J**), and a reduced swing speed in the back paws (**Fig. 3K**), indicating a compromised motor coordination. In the front paws, *Elp1*^cKO^ animals exhibit an increased step cycle duration (stance and swing phase of the paw), increased swing phase and decreased swing speed (**Supplemental Fig. S4A-C**). Most significant phenotypes were already present at week 8, and we did not observe any parameter worsening over time, suggesting that the cellular changes affecting the gait might take place at an earlier time point.

### Cerebella of adult Elp1^cKO^ ataxia mice are smaller, have fewer granule cells and show Purkinje cell abnormalities

Next, we investigated the underlying cellular changes driving the onset of the ataxia phenotype in *Elp1*^cKO^ mice. Following the last behavioral measurement at the age of 12 weeks, the animals were sacrificed and the gross morphology of the brains was analyzed (**Fig. 4A**). *Elp1*^cKO^ brains weighed less compared with *Elp1*^WT^ littermate brains (**Fig. 4B**). To identify the reason for the weight difference, we quantified the surface areas of brain regions as an approximate for brain region size (**Fig. 4A**). While the surface area of the whole brain and the cortex remained unaltered, the cerebellum surface area was significantly reduced in *Elp1*^cKO^ mice of both sexes (**Fig. 4C-E**).

**Figure 4.**
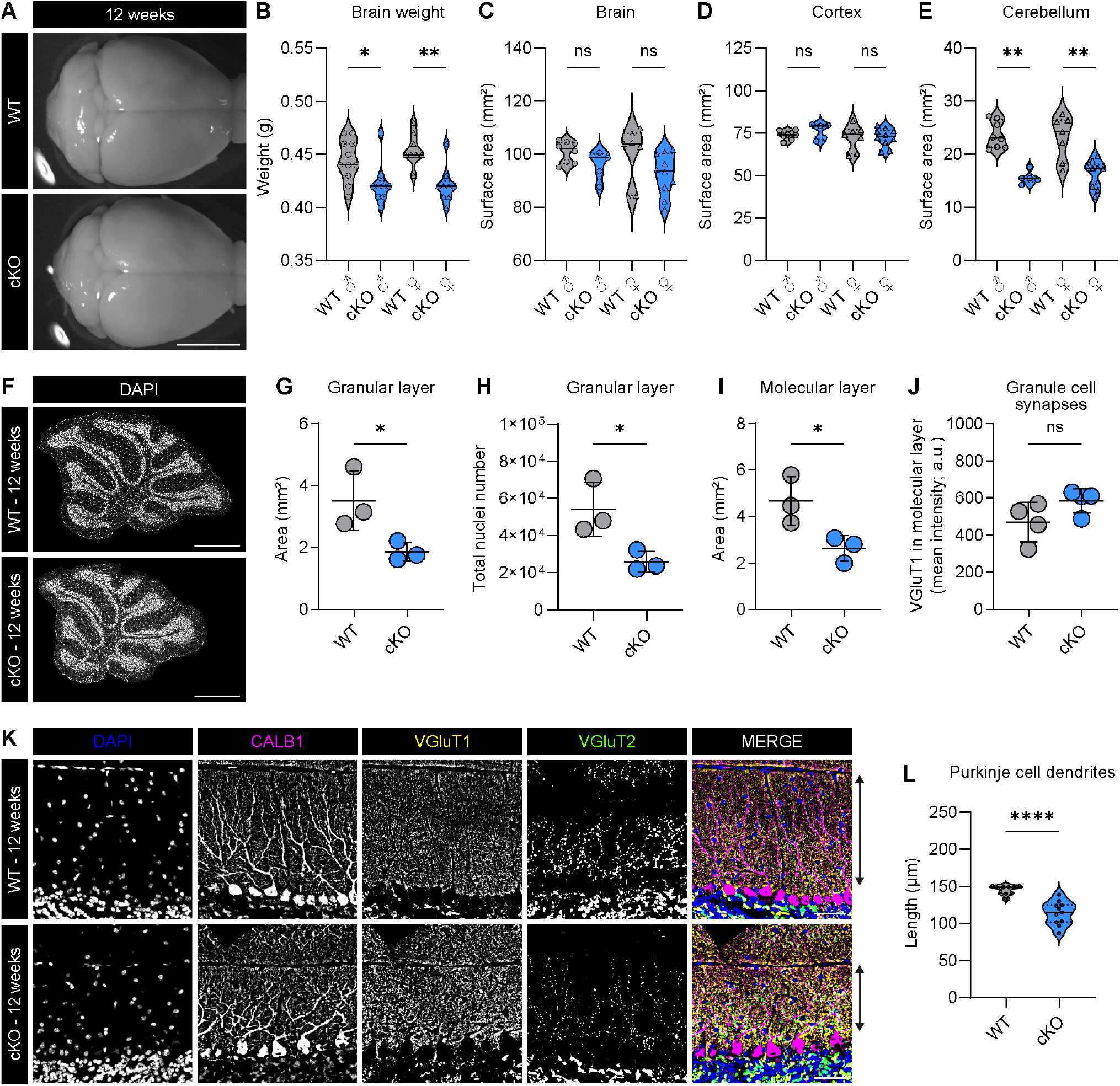
Ataxic, 12-week-old *Elp1*^cKO>^ mice have smaller cerebella, fewer GCs and a reduced area of the granular and molecular layers. (**A**) Representative brightfield images of brains of 12-week-old *Elp1*^WT^ and *Elp1*^cKO^ mice. Scale bar, 5 mm. (**B**) Total brain weight of 12-week-old *Elp1*^WT^ (gray) and *Elp1*^cKO^ (blue) mice. Mann-Whitney *U* test. *N* ≥ 8 brains per genotype and sex. (**C-E**) Quantification of surface areas from brightfield images: whole brain (**C**), cortex (**D**) and cerebellum (**E**). Mann-Whitney *U* tests. *N* = ≥ 5 brains per genotype and sex. (**F-I**) Representative nuclei (DAPI) staining of sagittal cerebellum section (**F**; scale bar, 1 mm) and quantification of granular layer area (**G**), total nuclei number in granular layer (**H**) and molecular layer area (**I**) in whole cerebellum overview image. Mean ± SD. Unpaired *t*-test. *N* = 3 (**J**) Quantification of VGluT1 mean fluorescence intensity in molecular layer of whole cerebellum. Mean ± SEM. Unpaired *t*-test. *N* = 4 (**K**) Representative immunofluorescence staining of DAPI (blue), CALB1 (magenta), VGluT1 (yellow) and VGluT2 (green) expression in lobe IV of 12-week-old *Elp1*^WT^ and *Elp1*^cKO^ cerebella. Arrow demonstrates width of molecular layer. Scale bar, 50 μm. (**L**) Quantification of maximum length of Purkinje cell dendrites. *N* = 3 mice/genotype, *n* = ≥ 3 sections/mouse.

To determine the underlying cause of the decreased cerebellar size in *Elp1*^cKO^ mice, we measured the area and total nuclei number of the granular layer, molecular layer, and PC layer. We found that the area of granular layer was reduced by about 47% in *Elp1*^cKO^, and the total number of nuclei in the granular layer in one section decreased by about 52% (**Fig. 4F-H**). The mean number of nuclei per mm^2^ remained unchanged (**Supplemental Fig. S5A**). We observed a decrease of the granular layer width in lobe III by 25% (**Supplemental Fig. S5B**), indicating that a decrease in width likely results in the reduced area of the granular layer. The strong decrease of granular layer area and nuclei suggests a diminished total number of GCs in *Elp1*^cKO^ cerebella. Next, we measured the molecular layer, where parallel fibers from the GCs innervate the dendritic trees of PCs. Similar to the granular layer, *Elp1*^cKO^ cerebella exhibit a reduced total area and width of the molecular layer (**Fig. 4I**; **Supplemental Fig. S5C**). We did not observe any differences in the width of the PC layer (**Supplemental Fig. S5D**).

Because GCs form excitatory VGluT1+ synapses with PCs via their parallel fibers in the molecular layer (Leto et al. (2016)), we hypothesized that the reduced pool of GCs would result in a decreased amount of VGluT1+ synapses. However, we did not observe any significant differences in VGluT1+ fluorescence intensity at week 12 in the molecular layer (**Fig. 4J,K**; **Supplemental Fig. S5E-F**). We also did not observe any differences in the density of climbing fiber synapses (VGluT2+) in the molecular layer and mossy fiber synapses (VGluT1+ and/or VGluT2+) in the granular layer (**Fig. 4K; Supplemental Fig. S5F-I**).

Pathologic changes during development and/or loss of PCs have been shown to cause cerebellar ataxias (Leto et al. 2016). To determine whether the ataxia phenotype might be evoked by PC loss, we quantified the number of PCs in different lobes. We did not observe any significant differences in the overall morphology or density of PCs between *Elp1*^cKO^ and *Elp1*^WT^ animals (**Fig. 4K, Supplemental Fig. S5F,J**). Similar to the reduced width of the molecular layer, the length of *Elp1*^cKO^ PC dendrites was also reduced (**Fig. 4K,L**).

Taken together, 12-week-old *Elp1*^cKO^ animals had smaller cerebella, most likely caused by a reduction of granular and molecular layers via a decrease in the total GC number in *Elp1*^cKO^ cerebella. Notably, we did not observe any differences in the PC numbers but dendritic complexity of PCs might be compromised due to shorter dendrites and reduced GC synaptic inputs, suggesting abnormal PC development as a result of defects of GC development.

### Early postnatal Elp1^cKO^ mice have smaller cerebella

Because total GC numbers were the only major defect we identified at week 12, we next examined animals prior to the onset of ataxia to determine when GC numbers were reduced. Similar to 12-week-old mice, P28 *Elp1*^cKO^ mice had smaller cerebella with a reduced area and fewer nuclei in the granular layer (**Supplemental Fig. S6A-D**), whereas no changes were detected to the molecular layer (**Supplemental Fig. S6A,E**).

Next, we examined the effect of *Elp1* loss to the GC lineage at P7, when GCP proliferation in the EGL peaks (Consalez et al. 2021). First, we assessed the motor control and coordination of P7 pups by testing the righting reflex (Feather-Schussler and Ferguson 2016); by P5, mouse pups are able to flip from a supine position to their paws (Heyser 2003). *Elp1*^cKO^ pups had proper righting reflexes (**Supplemental Fig. S7A**), suggesting no motor defects at this age. Next, we examined the total brain size and cellular composition of *Elp1*^cKO^ animals. We observed a reduced total weight of *Elp1*^cKO^ brains compared with their control littermates (**Fig. 5A-B**). While the total surface areas of brain, cortex and cerebellum did not change (**Fig. 5C,D**; **Supplemental Fig. S7B**), the ratio of cerebellum/cortex area was significantly reduced in *Elp1*^cKO^ mice (**Fig. 5E**), suggesting slightly smaller cerebella already at P7.

**Figure 5.**
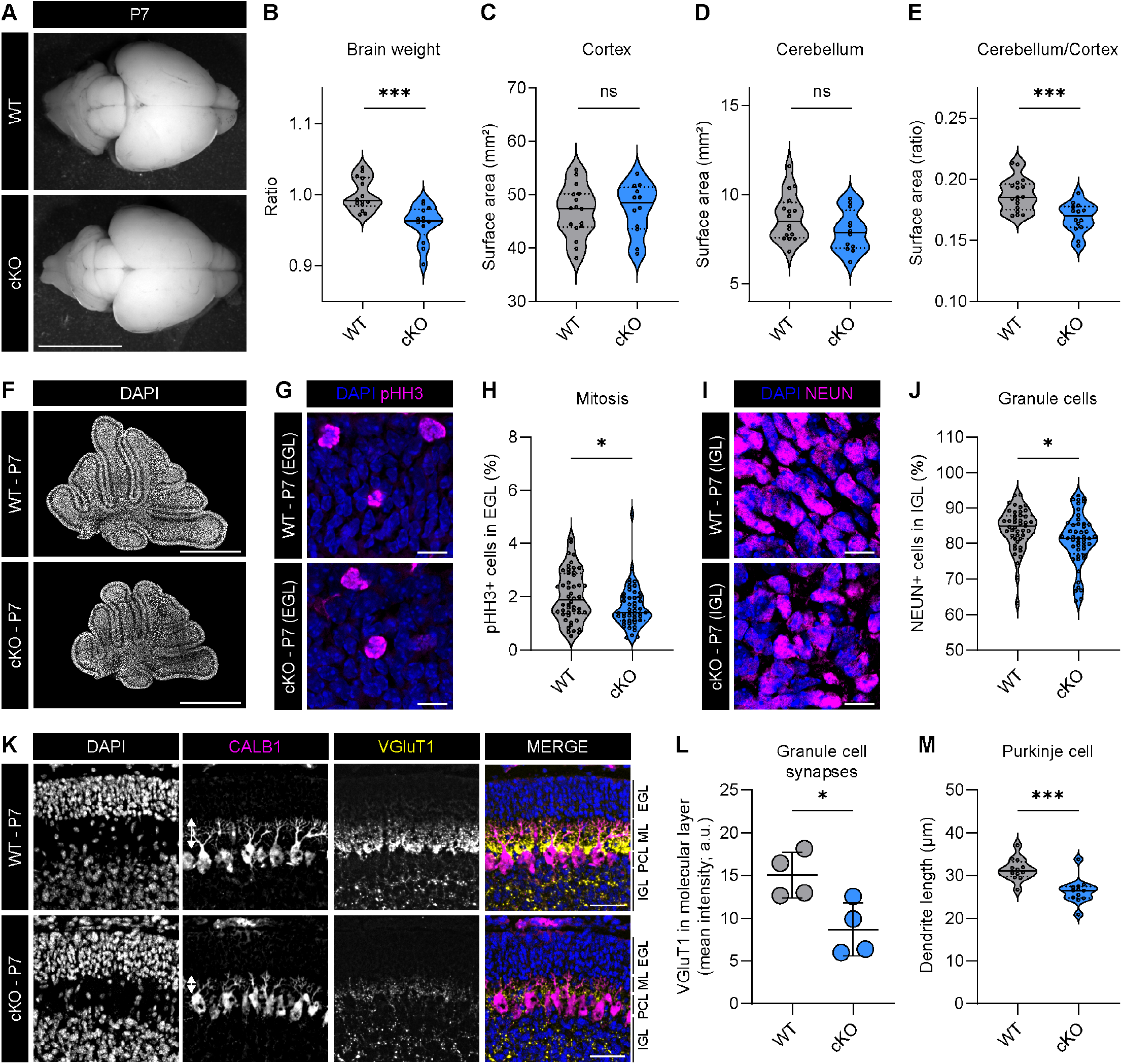
P7 *Elp1*^cKO^ mice show delayed GC maturation, resulting in reduced cerebellar size. (**A**) Representative brightfield images of *Elp1*^WT^ and *Elp1*^cKO^ brains at P7. Scale bar, 5 mm. (**B**) Quantification of P7 *Elp1*^cKO^ and *Elp1*^WT^ brain weights. Mann-Whitney *U* test. *N* = 14 (WT) and 13 (cKO). (**C-E**) Quantification of surface areas from brightfield images: cortex (**C**), cerebellum (**D**) and ratio of cerebellum/cortex (**E**). Unpaired *t-*tests. *N* = 16 (WT) and 12 (cKO). (**F**) Representative nuclei staining (DAPI) of sagittal section of whole cerebella. Scale bar, 1 mm. (**G,H**) Representative immunofluorescence staining (**G**) and quantification of DAPI (blue) and mitosis marker pHH3 (magenta; **H**) in EGL of lobes III, VI, IX and X at P7. Scale bar, 10 μm. *N* = 3 mice/genotype, *n* = 4 sections/lobe. (**I,J**) Representative immunofluorescence staining (**I**) and quantification (**J**) of DAPI (blue) and granule cell marker NEUN (magenta) in IGL of lobes III, VI, IX and X at P7. Scale bar, 10 μm. *N* = 3 mice/genotype, *n* = 4 sections/lobe. (**K**) Representative immunofluorescence staining of DAPI (blue), Purkinje cell marker CALB1 (magenta) and granule cell synapse marker VGluT1 (yellow) at P7. Scale bar, 50 μm. Arrows in CALB1 panel indicate length of Purkinje cell dendrites. IGL, internal granular layer. PCL, Purkinje cell layer. ML, molecular layer. EGL, external granular layer. (**L**) Quantification of VGluT1 mean fluorescence intensity in molecular layer of whole cerebellum at P7. Mean ± SD. Unpaired *t*-test. *N* = 4 mice/genotype. (**M**) Quantification of Purkinje cell dendrite length in lobe VI. Mann-Whitney *U* test. *N* = 3 mice/genotype, *n* = 4 sections/mouse.

At the cellular level, immunofluorescence stainings revealed no significant differences in the area or nuclei number of the EGL, IGL or ML (**Supplemental Fig. S7C-G**). In the EGL, the number of GCPs (PAX6+) was unchanged (**Supplemental Fig. S7H,I**) but slightly fewer mitotic GCPs (pHH3+) were present in *Elp1*^cKO^ cerebella (**Fig. 5G,H**). However, no significant differences in the number of cycling (MKI67+), S-phase (EdU+) or cell cycle exiting GCPs (EdU+MKI67-/EdU+) were detected (**Supplemental Fig. S7J-M**), which corresponds to the *in vitro* assay results above (**Fig. 2H**).

Upon differentiation, GCPs migrate inwards to the IGL and become mature NEUN+ GCs. Immunofluorescence imaging revealed a significant decrease of NEUN+ GCs in *Elp1*^cKO^ IGL (**Fig. 5I,J**). In addition, *Elp1*^cKO^ cerebella expressed significantly lower VGluT1 levels in the molecular layer (**Fig. 5K,L**; **Supplemental Fig. S8A**), further confirming fewer mature GCs in *Elp1*^cKO^ mice at P7. Furthermore, a lower density of VGluT1+ mossy fiber synapses were present in the IGL, whereas VGluT2+ synapses (mossy fiber) in the IGL and VGluT2+ synapses in both IGL (mossy fiber) and molecular layer (climbing fiber) were unaffected (**Fig. 5K**; **Supplemental Fig. S8A-D**). Total PC numbers were unchanged; instead, dendrite length was reduced in *Elp1*^cKO^ animals (**Fig. 5K,M**). Taken together, P7 *Elp1*^cKO^ cerebella display a delay in GC maturation, as the total number of GCPs in the EGL are unchanged, while the number of mature GCs are decreased. This delay is likely one underlying cause of the reduction in cerebellum size.

### Elp1^cKO^ increases cell cycle inhibition, apoptosis and reduces

#### differentiation

ELP1 is the scaffolding subunit of the Elongator complex, which catalyzes the addition of 5-carboxymethyl derivatives to tRNAs and modulates protein expression during translation. Previous work has shown that *ELP1* loss can lead to various outcomes such as protein mis-folding and aggregation, the activation of the unfolded protein response, codon-dependent translational reprogramming and/or the disruption of protein homeostasis (Laguesse et al. 2015; Nedialkova and Leidel 2015; Goffena et al. 2018; Waszak et al. 2020). Therefore, we measured Elongator-dependent tRNA modifications (mcm^5^, ncm^5^ and mcm^5^s^2^) from *Elp1*^cKO^ and *Elp1*^WT^ bulk GCPs isolated from P7 cerebellum. However, we did not detect changes in these modifications (**Supplemental Fig. S9A-C**), likely because of their low abundance in bulk tRNA and/or the incomplete *Elp1* knockout in GCPs in the nodular lobes IX or X (**Fig. 2B**).

To identify the molecular causes underlying the delay in GC maturation and reduced mature GC number, we performed bulk RNA-sequencing of P7 *Elp1*^WT^ and *Elp1*^cKO^ GCPs (**Fig. 6A**; **Supplemental Table S5**). We found that *VGluT1* expression was strongly reduced in *Elp1*^cKO^ GCPs, providing a molecular basis for reduced protein expression in mature GCs at P7 (**Fig. 5K,L**). Furthermore, cellular stress genes *Asns* and *Sesn2*, the pro-apoptotic factor *Pmaip1* (also called *Noxa*) and the unfolded protein response pathway member *Chac1* showed a significant upregulation in *Elp1*^cKO^ GCPs (**Fig. 6A**). In addition, cell cycle inhibitor *Cdkn1a* (or *P21*) expression was highly upregulated in *Elp1*^cKO^ GCPs. Confirming the individual genes we identified as mis-regulated, Gene Set Enrichment Analysis (GSEA, Subramanian et al. 2005) of the transcriptomic data revealed significant upregulation of P53-mediated apoptosis pathway and downregulation of neuronal development and synaptic transmission pathways (**Fig. 6B: Supplemental Fig. S9D**), indicating a compromised neuronal maturation in *Elp1*^cKO^ GCPs. Altogether, *Elp1*^cKO^ GCP transcriptomes have elevated expression of cell cycle inhibitors, stress sensors and pro-apoptotic factors, suggesting an increased intrinsic cell stress in *Elp1*^cKO^ GCPs.

**Figure 6.**
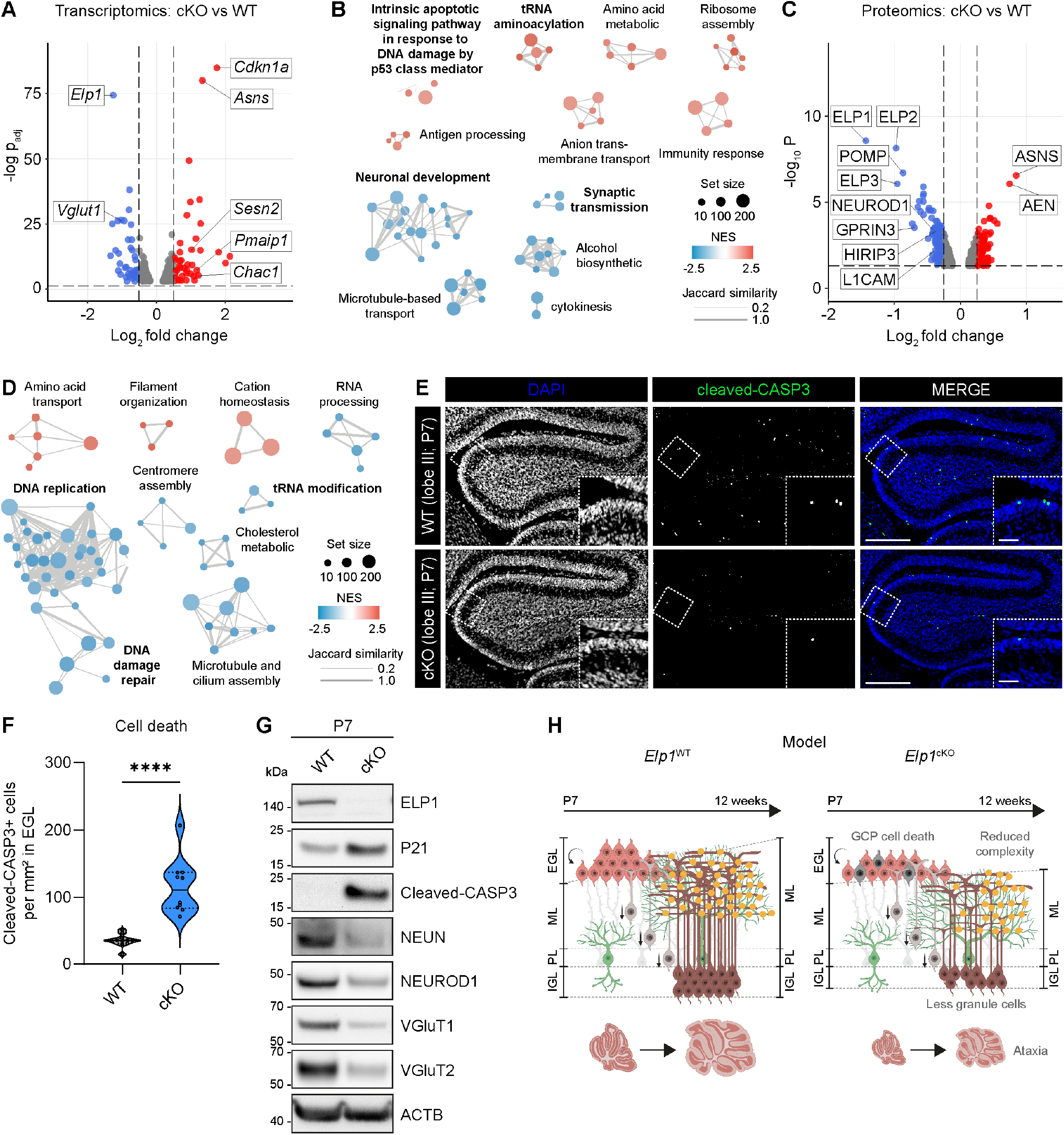
*Elp1*^cKO^ increases cell cycle inhibition and cell death in GCPs and reduces GC differentiation. (**A**) Volcano plot of bulk RNA-seq of P7 *Elp1*^cKO^ vs *Elp1*^WT^ GCPs revealing up- (red) and downregulated (blue) genes/proteins. *N* = 3 samples/genotype. Raw data is available in Supplemental Table S5. (**B**) Network visualization of GSEA of transcriptomes shows up- (red) and downregulated (blue) biological processes. Size of dot represents gene set size. *p*_adj_ < 0.05. NES, normalized enrichment score. *N* = 4 samples/genotype, matched with proteomics samples. (**C**) Volcano plot of proteomics, highlighting up- (red) and downregulated (blue) proteins in P7 *Elp1*^cKO^ vs *Elp1*^WT^ GCPs. *N* = 4 samples/genotype. (**D**) Proteome network analysis revealing up- (red) and downregulated (blue) biological processes. *p*_adj_ < 0.05. (**E**) Representative immunofluorescence staining of DAPI (blue) and cell death marker cleaved-CASP3 (green) in lobe III. Single z-slice. Scale bar, 250 μm. Inset, scale bar, 50 μm. (**F**) Quantification of cleaved-CASP3 positive cells in external granular layer (EGL) of whole cerebellum. Mann-Whitney *U* test. *N* = 3 mice/genotype. *n* = 2-4 sections. (**G**) Immunoblot validation of ELP1, P21, cleaved-CASP3, NEUN, NEUROD1, VGluT1, VGluT2 and ACTB protein abundance in lysates from *Elp1*^WT^ and *Elp1*^cKO^ GCPs at P7. (**H**) Schematic model of ataxia onset by *Elp1*^cKO^ in GCPs, presumably through GCP cell death in early stages, resulting in a reduced GC pool, decreased GC-PC synaptic complexity and smaller cerebellum size.

To examine changes at the protein level, we performed Tandem mass tag-mass spectrometry proteomics using P7 GCPs. This method validated the significant reduction of ELP1, ELP2, and ELP3 protein levels, confirming the destabilization of the Elongator complex (**Fig. 6C**; **Supplemental Fig. S9E**; **Supplemental Table S5**) as observed by immunoblotting before (**Fig. 2C-G**). Further, neurite outgrowth factors HIRIP3, GPRIN3 and L1CAM and GC differentiation factor NEUROD1 were reduced, suggesting the molecular reasons of impaired neuronal differentiation in *Elp1*^cKO^ GCPs (**Fig. 6C**; **Supplemental Fig. S9E**). Upregulation of stress sensor ASNS and apoptosis-enhancing nuclease AEN suggested increased cell death in *Elp1*^cKO^ GCPs. Furthermore, proteomic GSEA revealed the significant downregulation of biological processes associated with DNA replication and DNA damage repair (**Fig. 6D**), indicating decreased cell cycle activity.

To verify that *Elp1*^cKO^ GCPs were undergoing cell stress leading to apoptotic cell death, we performed immunofluorescence staining against cleaved-CASP3, the functional unit of the cell death effector gene *Caspase3*. Indeed, we confirmed a marked increase of dying GCPs in the EGL of *Elp1*^cKO^ mice compared with *Elp1*^WT^ sibling controls (**Fig. 6E,F**). The increased expression of cleaved-CASP3 was further confirmed by immunoblotting of purified GCPs (21-fold increase; **Fig. 6G**; **Supplemental Fig. S9F**). We also confirmed increased P21 protein levels, with a 5-fold increase in *Elp1*^cKO^ GCPs (**Fig. 6G; Supplemental Fig. S9G**). Overall, the combined increase of P21 and cleaved-CASP3 in *Elp1*^cKO^ GCPs suggests activation of cell cycle inhibition resulting in cell death, ultimately leading to a reduced number of GCPs and subsequent mature GC pool. Reduced GCP differentiation was also confirmed by lower level of mature GC markers NEUN, NEUROD1 and VGluT1 (**Fig. 6G**; **Supplemental Fig. S9H-K**).

Collectively, *Elp1* loss in GCPs increased cell cycle inhibition and cell death pathways and decreased terminal differentiation into GCs. Both processes likely reduce the final pool of functional GCs and, consequently, functional synapses with PCs, eventually leading to the gradual onset of ataxia (**Fig. 6H**).

## DISCUSSION

### Role of granule cells during motor behavior and ataxia

Using an *Atoh1-Cre* mouse line, we removed *Elp1* in the majority of the cerebellar GCPs. Adult mice developed ataxia, which we propose is the result of increased cell death of GCPs at early postnatal stages leading to decreased mature GCs as animals age. While pathologic changes or loss of PCs are well-established drivers of cerebellar ataxia (Becker et al. 2009; Hourez et al. 2011; Todorov et al. 2012; Tsai et al. 2012; Hansen et al. 2013), manipulation of GCs have also been shown to impair motor coordination and evoke signs of cerebellar ataxia (Yamamoto et al. 2003; Wada et al. 2007; Kim et al. 2009; Galliano et al. 2013; Miyazaki et al. 2021; Lee et al. 2023). In our model, the defects in both GCs and PCs in adult *Elp1*^cKO^ animals were subtle, although the ataxia phenotype was strong. In recent studies, GCs were either selectively ablated or functionally silenced, causing severe motor deficits affecting balance, coordination and gait (Miyazaki et al. 2021; Lee et al. 2023). While selective GC ablation resulted in the reduction of cerebellum size and the width of both GL and ML, PC death was not apparent and also the overall PC firing rates were unaffected (Miyazaki et al. 2021; Lee et al. 2023). Correspondingly, *Elp1*^cKO^ PC numbers were unchanged and only the length of the dendrites was reduced, which might be caused by the reduction of molecular layer width. Together with our results, recent studies suggest that reduced GC signaling by depletion of synaptic transmission or by reduction of the total GC pool results in motor deficits, while leaving PC function mostly intact (Miyazaki et al. 2021; Lee et al. 2023).

### ELP1 function in granule cells: a potential link to Familial Dysautonomia?

FD patients typically present with progressive ataxias (Portnoy et al. 2018) although the underlying cause of the ataxia is unclear. In FD patients, loss of muscle spindle afferents in the peripheral nervous system plays a part in the ataxic gait (Macefield et al. 2011). While white matter loss has been observed in the cerebellar peduncles, the cerebellar vermis of FD patients does not have an obvious reduction in size (Axelrod et al. 2010). In older patients, however, generalized atrophy may occur in the cerebellum (Cohen and Solomon 1955; Axelrod and Gold-von Simson 2007). Our studies demonstrate a central role of *Elp1* in maintaining GC (progenitor) health and a functional cerebellum. Given that tissue-specific splicing in the cerebellum of FD patients leads to an increase in mis-spliced *ELP1* gene (Hims et al. 2007) and the observation that cerebellar atrophy occurs with age (Axelrod and Gold-von Simson 2007), *ELP1* loss in the cerebellum of FD patients may indeed play a role, albeit likely minor, in maintaining balance and coordination. Future studies in FD patients and patient-derived cells would be required to confirm this putative role in the future.

### ELP1 and Elongator are crucial for cerebellar neurodevelopment and maintenance

In mouse models, homozygous loss of *Elp1* is embryonically lethal by E12.5, with a strong developmental delay, heart and brain defects, and incomplete neurulation (Chen et al. 2009; Dietrich et al. 2011). CNS-specific knockout of *Elp1* using a *Tuba1a-Cre* mouse line resulted in reduced ELP1 expression in the striatum and cortex (Chaverra et al. 2017). While the effect on the cerebellum was not explicitly examined in this mouse model, animals had a slow and unsteady gait. In this study, *Elp1* was critical for cell survival of spinal motor neurons and cortical neurons (Chaverra et al. 2017). Similarly, knockout of *Elp1* in neural crest-derived neurons resulted in increased neuronal cell death of selective populations and in diminished nerve growth as development proceeded (George et al. 2013; Jackson et al. 2014; Leonard et al. 2022). Overall, combining these published studies with our current work confirms the essentiality of *Elp1* in maintaining neuronal survival as a key function in maintaining brain health. Whether this neuronal cell death results from the direct or indirect effect of impaired tRNA modifications is still unknown (Ueki et al. 2018); it is clear, however, that only specific classes of neurons are sensitive to *Elp1* loss. Future work is required to determine the underlying mechanisms of this sensitivity.

Other members of the Elongator complex have also been implicated in other neurodevelopmental disorders, including intellectual disability and autism spectrum disorder (Addis et al. 2015; Kojic et al. 2021), suggesting a central role of this complex in CNS development. Specific patientderived germline mutations of *Elp2* in mice cause gait defects, although specific defects in the cerebellum were not identified (Kojic et al. 2021). Mice with a germline mutation in *ELP6*, however, develop ataxia, likely as a result of PC degeneration (Kojic et al. 2018). Altogether, the Elongator complex plays a crucial role in developing and maintaining health of cerebellar neurons.

## Author Contributions

F.M., P.B.G.d.S., M.E.S., J.G.-L., S.T.A., L.Sp., L.Si., M.Ra., M.Ro. and L.M.K. performed experiments. F.M., M.E.S., and L.B. performed image quantification. Y.L., P.J., F.M. and M.S. analyzed proteomics and transcriptomic data. C.L. performed the behavior studies and F.M., A.P., and L.M.K. analyzed the behavioral data. M.Z., H.L., P.A.N., and A.P. advised on the experimental design. L.M.K. designed and supervised the study. F.M. and L.M.K. wrote the manuscript. All authors read and approve the final version of the manuscript.

## Acknowledgments

This study was supported by a DFG Individual Research Grant to L.M.K. (#497317859). We thank the following core facilities at the DKFZ: High-Throughput Sequencing unit of the Genomics & Proteomics Core Facility, the Light Microscopy Core Facility, the Flow Cytometry Core Facility and the Omics IT and Data Management Core Facility. We thank Claudia Pitzer and Barbara Kurpiers at the INBC at University of Heidelberg, the Imaging facilities at EMBL, and Marie-Claire Indilewitsch for genotyping. Cartoon panels created with biorender.com.

## Materials and Methods

### Animals

*Elp1* floxed mice were generated by breeding *Elp1* knockout first animals (*Elp1*^tm1a(KOMP)Wtsi^; George et al. (2013)) to ubiquitous *Flp* expressing mice (Rodríguez et al. (2000); JAX, #005703). To generate *Elp1* conditional knockout mice, *Elp1*^fl/fl^ animals were crossed with *Atoh1-Cre* (Matei et al. (2005); JAX, #011104). Genotyping primer sequences available in Supplemental Table S1. Mice were bred on a C57BL/6N background. Animals were housed in a temperature-controlled vivarium on a 12:12h light-dark cycle with *ad libitum* food and water. Animals of both sexes were used for experiments. For controls, we used Cre-negative littermates. All animal experiments for this study were conducted according to the animal welfare regulations approved by the responsible authorities in Baden-Württemberg, Germany (Regierungspraesidium Karlsruhe, approval numbers: G-172/18, G-136/23).

### Motor behavior studies

Motor behavior studies were performed at the Interdisciplinary Neurobehavioral Core (INBC) at the University of Heidelberg. Data collection was performed blinded to genotype. Experiments were carried out during the light phase. Animals were acclimated to the test room for at least 1-2 days before testing. Weight measurements, rotarod, grip strength, and CatWalk tests were performed at 8, 10, and 12 weeks of age. Raw data is available in Supplemental Table S2-4.

### SHIRPA, Rotarod, Grip Strength assay

A modified SHIRPA Primary Screen (Rogers et al. 1997) was performed visually one time at week 7 and served as a general acclimation phase and as an assessment of overall animal health. General behavioral and functional parameters were measured by a ranking score: gait, body tone, motor control and coordination, aggression, salivation, lacrimation, piloerection, defecation and temperature. Rotarod and grip strength tests were performed as described in Eltokhi et al. (2021). Briefly, we used an accelerating Rotarod system, and tested animals for a maximum of 8 minutes. Animals completed 3 trials per day with a break (10 minutes) between the trials, and measurements were averaged. For the grip strength assay, animals pulled on a fixed stirrup and the strength of the front paw was measured three times; measurements were averaged.

### Catwalk test

For gait analysis, a digital Catwalk was used (CatWalk XT, Noldus Information Technology BV, Hamers et al. (2006)). Animals were placed at one end of a 130-cm tube, and the parameters were automatically calculated. Animals were tested three times per time point and measurements were averaged. Raw data is available in Supplemental Tables S3 and S4.

### Righting reflex

Righting reflex was tested at P7. Animals were placed in the supine position and held in this position for 5 seconds. After, the animals were released and the time taken for the animal to return to the prone position was recorded. The experiment lasted a maximum of one minute. The test was repeated a total of three times.

### EdU injections

P7 mice of the appropriate genotype were injected intraperitoneally with 10 mg/ml stock solution of EdU (50 mg/kg, 20 μl total volume, dissolved in PBS, Invitrogen). Mice were sacrificed at 2h post-injection. Brains were harvested, fixed overnight in 4% paraformaldehyde (PFA) at 4°C, soaked in 30% sucrose in PBS overnight, and embedded in Optimal Cutting Temperature (OCT) compound. Frozen 10 μm-thick sections were collected on Fisher Superfrost Plus slides using a cryostat and treated with Click-iT™ EdU imaging reagent (Invitrogen).

### Tissue preparation, immunofluorescence staining, and quantification

Adult mice were anaesthetized with a ketamine/xylazine/ acepromacin mix, transcardially perfused with 4% PFA in phosphate-buffered saline (PBS, pH 7.4). After, brains were harvested, post-fixed in 4% PFA at 4°C, soaked in 30% sucrose in PBS overnight, and embedded in OCT compound. Immunofluorescence staining was performed as previously described (Kutscher et al. 2020). Frozen 10 μm-thick sections were collected on Fisher Superfrost Plus slides using a cryostat. Sections were blocked for 30 min at RT with 10% normal donkey serum (NDS) and incubated with primary antibody overnight at 4°C. After washing with PBST, sections were incubated with appropriate fluorescence secondary antibodies for 1h at RT. Slides were mounted in ProLong Gold Mountant (Invitrogen). Antibodies used are listed in Supplemental Table S1. All slides were counterstained with DAPI (300 nM). Imaging and quantification were performed blinded to genotype. For display, comparative immunostainings were edited with identical settings in FIJI (Schindelin et al. 2012). For quantification, the number of average positive cells was counted in 3-4 100 μm x 100 μm regions in 3 independent cerebella in the same lobe and region, as indicated in the figures. To quantify areas or mean fluorescence intensities from microscopy images, brain regions or layers were segmented manually and measured in FIJI.

### GCP isolation, culture and proliferation assay

GCP isolation was performed as previously described (Hatten 1985; Kawauchi et al. 2012) and cell density was diluted to 2.5x10^6^ cells/ml with GCP media. For *in vitro* studies, GCPs were plated on 6-well (2 ml, immunoblot) and 24-well plates (0.4 ml; qPCR or EdU proliferation assay) for 72 h in the presence or absence of 200 nM Smoothened agonist (Merck). Full medium changes were performed daily. Cell proliferation assay was performed as described in Haag et al. (2021). Briefly, a final concentration of 10 μM EdU was added to the culture medium and incubated for 1 h. Proliferative cells were labeled using the Click-iT EdU Alexa Fluor 647 Flow Cytometry Assay Kit (Life Technologies) according to manufacturer’s instruction.

### Quantitative PCR (qPCR)

RNA was extracted with the Maxwell® RSC simplyRNA tissue kit (Promega) and cDNA was generated with SuperScript II kit (Invitrogen). qPCR was performed with SYBR green PCR master mix (Applied Biosystems) with 10 ng cDNA input in triplicates on a QuantStudio 5 real-time PCR system (Thermo Fisher Scientific). Primer sequences are listed in Supplemental Table S1. Relative gene expression was compared to reference gene *Tbp1*.

### RNA extraction and RNA-seq analysis

Tissue was homogenized using a QIAshredder (Qiagen). RNA was extracted with RNeasy Mini Plus Kit (Qiagen) following supplier’s instructions. For measurement of RNA quantity, ND1000 Spectrophotometer (NanoDrop Technologies) and/or Agilent 2100 Bioanalyzer was used.

FastQC-v0.11.8 was used for quality control of RNA-seq reads. BBDuk (minlen=30; others default) was used to remove remaining adapters and low-quality bases. The alignment of raw FASTQs was performed using STAR v2.5.3a tool (Dobin et al. 2013) to mouse reference genome (mm10). Count matrices were analyzed for differentially expressed genes with R package DESeq2 (Love et al. 2014). GSEA was performed with the R package clusterProfiler (v4.6; Wu et al. (2021)) based on the gene ontology biological processes gene set from MSigDB (v7.4.1) with a *p*_adj_ < 0.05 (Subramanian et al. 2005; Liberzon et al. 2011). GSEA results were visualized in Cytoscape with network edges defined by Jaccard distance between gene set pairs.

### Analysis of snRNA-seq data

To profile *ELP1*/*Elp1* expression in cerebellar cell types, we used processed snRNA-seq data covering cerebellum development in human and mouse (Sepp et al. 2024) as available at https://apps.kaessmannlab.org/sc-cerebellumtranscriptome/. The data was filtered for the main cerebellar cell types (VZ_neuroblast, parabrachial, GABA_DN, Purkinje, interneuron, NTZ_neuroblast, glut_DN, isth_N, GC/UBC, GC, UBC, astroglia, oligo, ependymal, meningeal, immune), and grouped by biological replicate and cell type, requiring at least 50 cells per group. Two values were calculated for each group: fraction of cells positive (n[UMI] 1) for *ELP1* and mean expression per cell (*ELP1* n[UMI] / total n[UMI]). These values were then standardized separately for the data produced with the two Chromium versions (v2 and v3; 10x Genomics), using version-specific means and standard deviations calculated based on values averaged across the developmental stages and cell types that were covered by both Chromium versions. The obtained z-scored data were averaged across groups for presentation; in case of the cell fraction the values were back-transformed using the mean and standard deviation values averaged across the versions.

### Protein extraction and immunoblotting

Protein extraction and immunoblotting was performed as described in Haag et al. (2021). Membranes were stripped and reprobed. Antibodies and reagents are listed in Supplemental Table S1.

### Protein Digestion and Tandem-Mass-Tag (TMT) Labeling

The analysis was performed with a previously optimized protocol (Pagala et al. 2015; Bai et al. 2017). For whole proteome profiling, quantified protein samples (300 μg in the lysis buffer with 8 M urea) for each TMT channel were proteolyzed with Lys-C (Wako, 1:100 w/w) at 21ºC for 2 h, diluted by 4-fold to reduce urea to 2 M for the addition of trypsin (Promega, 1:50 w/w) to continue the digestion at 21ºC overnight. The insoluble debris was kept in the lysates for the recovery of insoluble proteins. The digestion was terminated by the addition of 1% trifluoroacetic acid. After centrifugation, the supernatant was desalted with the Sep-Pak C18 cartridge (Waters), and then dried by Speedvac (Thermo Fisher). Each sample was resuspended in 50 mM HEPES (pH 8.5) for TMT labeling and then mixed equally, followed by desalting for the subsequent fractionation. 0.1 mg protein per sample was used.

### Extensive Two-Dimensional Liquid Chromatography-Tandem Mass Spectrometry (LC/LC-MS/MS)

The TMT labeled samples were fractionated by offline basic pH reverse phase LC, and each of these fractions was analyzed by the acidic pH reverse phase LC-MS/MS (Xu et al. 2009; Wang et al. 2014). We performed a 160 minute offline LC run at a flow rate of 400 μl per minute on an XBridge C18 column (3.5 μm particle size, 4.6 mm x 25 cm, Waters; buffer A: 10 mM ammonium formate, pH 8.0; buffer B: 95% acetonitrile, 10 mM ammonium formate, pH 8.0; Bai et al. (2017)). A total of 80 two-minute fractions was collected. Every forty-first fraction was concatenated into 40 pooled fractions which were subsequently used for whole proteome TMT analysis.

In the acidic pH LC-MS/MS analysis, each fraction from basic pH LC was dried by a Speedvac and was run sequentially on a column (75 μm x 35 cm for the whole proteome, 50 μm x 30 cm for whole proteome, 1.9 μm C18 resin from Dr. Maisch GmbH, 65ºC to reduce backpressure) interfaced with a Fusion MS (Thermo Fisher) for the whole proteome where peptides were eluted by a 90 min gradient (buffer A: 0.2% formic acid, 5% DMSO; buffer B: buffer A plus 65% acetonitrile). MS settings included the MS1 scan (410-1600 m/z, 60,000 resolution, 1 × 106 AGC and 50 ms maximal ion time) and 20 data-dependent MS2 scans (fixed first mass of 120 m/z, 60,000 resolution, 1 × 105 AGC, 200 ms maximal ion time, HCD, 36% normalized collision energy, 1.0 m/z isolation window with 0.2 m/z offset, and 20 s dynamic exclusion). MS settings included the MS1 scan (450-1600 m/z, 60,000 resolution, 1 × 106 AGC and 50 ms maximal ion time) and 20 data-dependent MS2 scans (fixed first mass of 120 m/z, 60,000 resolution, 1 × 105 AGC, 110 ms maximal ion time, HCD, 36% normalized collision energy, 1.0 m/z isolation window with 0.2 m/z offset, and 10 s dynamic exclusion).

### tRNA modification analysis

tRNA modification analysis was performed by Arraystar Inc. (Rockville, USA). Briefly, tRNAs were isolated from total RNA by Urea-PAGE electrophoresis. 60-90 nt tRNA bands were excised, purified by ethanol precipitation and quantified by Qubit RNA HS assay (Thermo Fisher). tRNAs were hydrolyzed to single nucleosides, dephosphorylated and deproteinized using 10.000-Da MWCO spin filter (Sartorius). Analysis of nucleoside mixtures was performed on Agilent 6460 QQQ mass spectrometer with an Agilent 1260 HPLC system. LC-MS data was acquired using Agilent Qualitative Analysis software. MRM peaks of each modified nucleoside were extracted and normalized to the sum of all modifications peak area.

### Statistical Analysis

Statistical analysis was performed with GraphPad Prism (version 10.2.0). Data was tested for normal distribution and parametric tests (unpaired *t*-test) were used only if the data was normally distributed. Otherwise, a nonparametric test as Mann-Whitney *U* test was chosen. Statistical significance between groups was considered when *P* values ≤ 0.05; significance levels are *P*<0.05 (^*^), *P*<0.005 (^**^), *P*<0.0005 (^***^), and *P*<0.0001 (^****^), if not otherwise indicated. The statistical test used for each analysis is listed in each figure legend. Data presentation, applied statistical tests, biological replicate number *N* and technical replicate number *n* are indicated in the figure legends.

## Data availability

All processed data is included in Supplemental Tables within the manuscript. Raw RNA-seq data is available upon request.

---

**Supplemental Figure S1.**
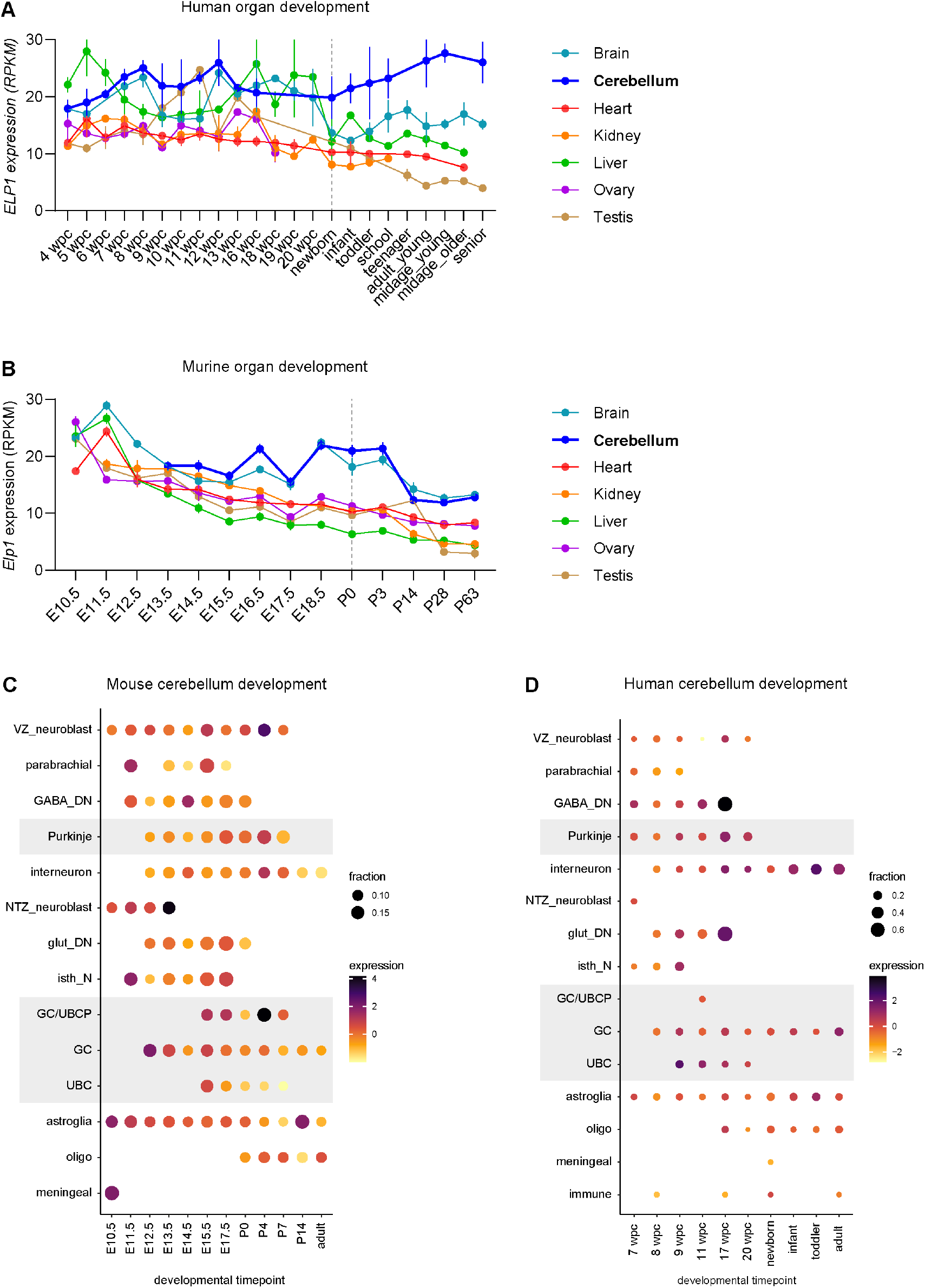
Human *ELP1* expression levels increase after birth and remain high in granule cells. (**A,B**) Bulk expression levels of human *ELP1* (**A**) and mouse *Elp1* (**B**) across different organs and developmental time points. Vertical dotted lines represent time point of birth. Expression data taken from Cardoso-Moreira et al. (2019). (**A**) *N* = 18-53 human donors per tissue. Line plot is adapted from Waszak et al. (2020). (**B**) *N* = 27-55 mouse samples per tissue. Mean ± SEM. RPKM, reads per kilobase of transcript per million mapped reads. wpc, weeks post conception. (**C,D**) Cell type-specific expression of mouse *Elp1* (**C**) and human *ELP1* (**D**) during cerebellar development based on single-nucleus RNA-sequencing atlases (Sepp et al. 2024). Dot size and color indicate fraction of cells and mean expression of *Elp1*/*ELP1* averaged across biological replicates, respectively. Areas with grey background are shown in Fig. 1A,B. GABA_DN, GABAergic deep nuclei neurons. GC/UBCP, granule cell/unipolar brush cell progenitor. glut_DN, glutamatergic deep nuclei neurons. GC, granule cell. isth_N, isthmic nuclei neurons. NTZ_neuroblast, neuroblasts migrating to the nuclear transitory zone. UBC, unipolar brush cell. VZ_neuroblast, neuroblasts originating from the ventricular zone.

**Supplemental Figure S2.**
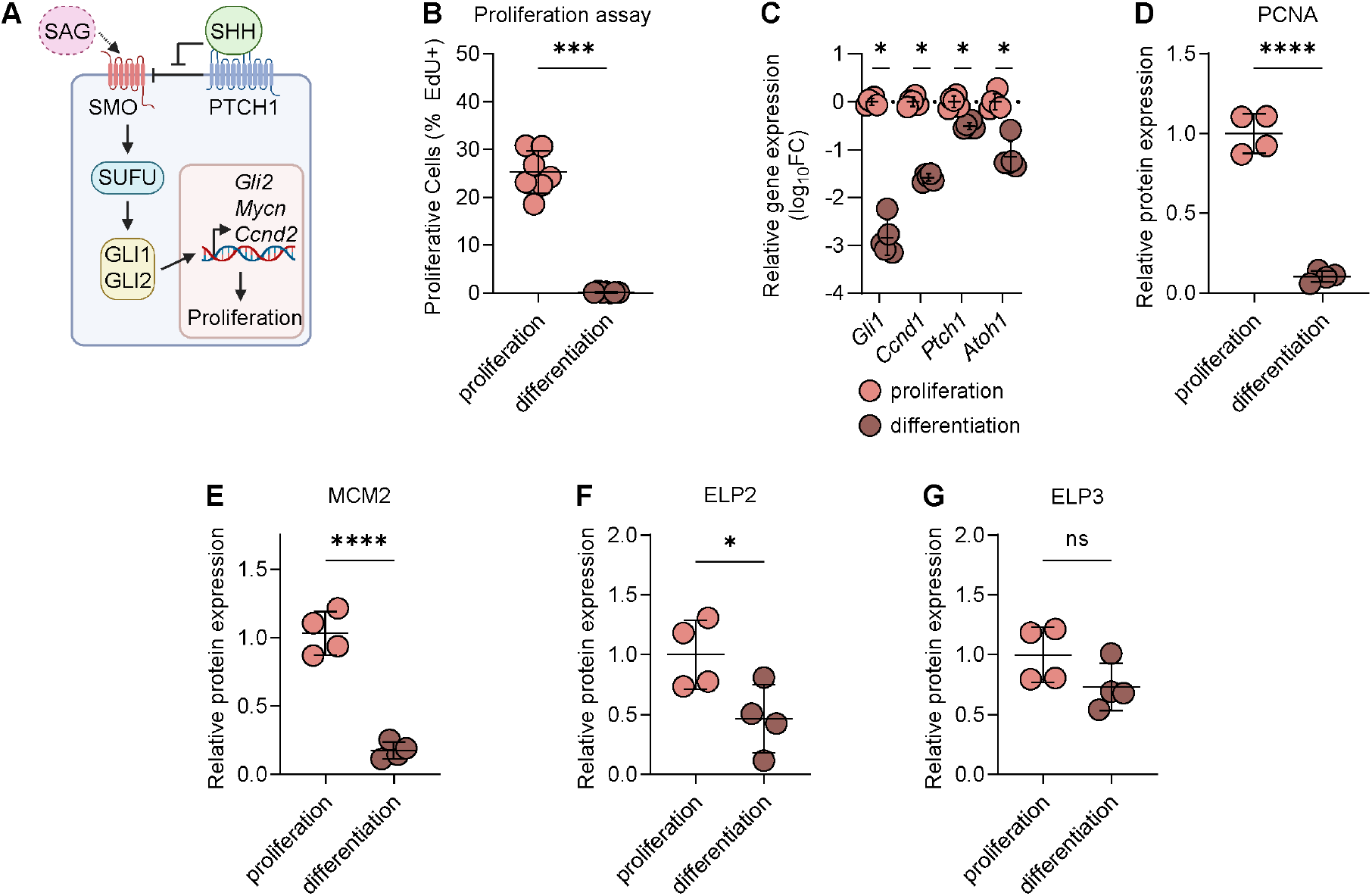
Increased ELP1 expression in proliferating GCPs. (**A**) Cartoon showing the SHH signaling pathway and target genes. SAG, Smoothened agonist. SHH, sonic hedgehog. (**B**) Percentage of proliferative GCPs (EdU+) after 72 h *in vitro* culture in the presence (orange, proliferation) or absence of SAG (brown, differentiation). Mann-Whitney *U* test. Mean ± SD. *N* = 7. (**C**) Relative mRNA expression of *Gli1, Ccnd1, Ptch1* and *Atoh1* (quantitative PCR) of GCPs cultured in the presence of SAG (orange, proliferation) or absence of SAG (brown, differentiation). Mann Whitney U test. Mean ± SD. *N* = 5. (**D-G**) Quantification of relative protein abundance of PCNA (**D**), MCM2 (**E**), ELP2 (**F**) and ELP3 (**G**) in GCP lysates normalized to ACTB. Unpaired *t*-tests. Mean ± SD. *N* = 4.

**Supplemental Figure S3.**
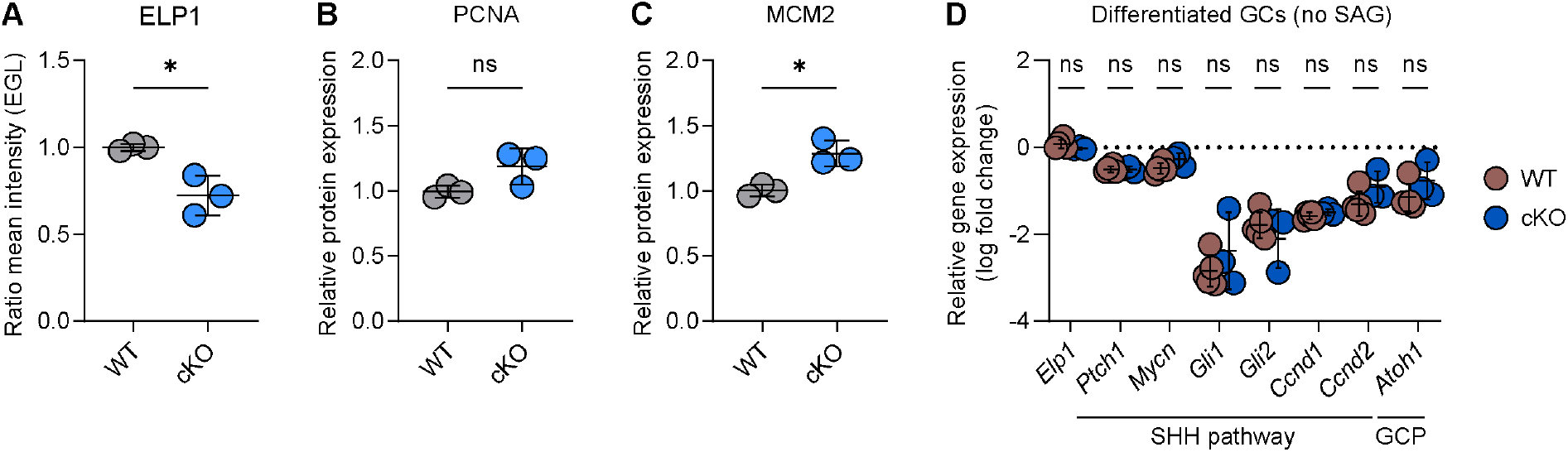
*Elp1*^cKO^ reduces ELP1 in the EGL but does not affect proliferation or response to SHH pathway stimulation. (**A**) Quantification of mean fluorescence intensity of ELP1 in the external granular layer (EGL) of whole cerebellum of P7 *Elp1*^WT^ and *Elp1*^cKO^ mice. Average mean fluorescence in *Elp1*^WT^ animals was used to calculate ratios. Unpaired *t*-test. Mean ± SD. *N* = 3. (**B,C**) Quantification of immunoblots against proliferation markers PCNA (**B**) and MCM2 (**C**) in *Elp1*^WT^ and *Elp1*^cKO^ GCP lysates, normalized to ACTB. Unpaired *t*-tests. Mean ± SD. *N* = 3. (**E**) Relative mRNA expression (quantitative PCR) of *Elp1*, SHH pathway genes (*Ptch1, Mycn, Gli1, Gli2, Ccnd1, Ccnd2*), and GCP marker *Atoh1* in differentiated, untreated *Elp1*^WT^ (dark brown) and *Elp1*^cKO^ (dark blue) GCs after 72 h *in vitro* culture. Mann Whitney U tests. Mean ± SD. *N* = ≥ 4.

**Supplemental Figure S4.**
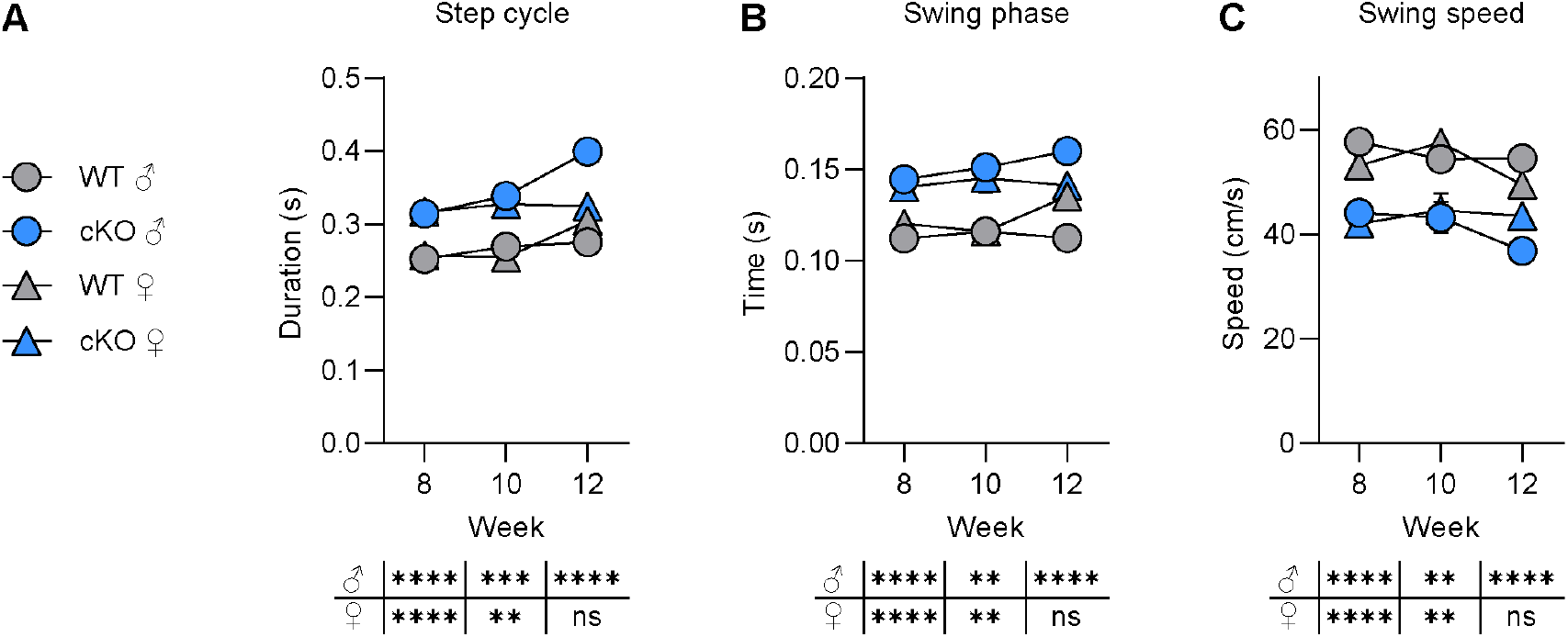
*Elp1*^cKO^ mice show compromised gait parameters in front paws. (**A-C**) Quantification of CatWalk parameters step cycle (**A**), swing phase (**B**) and swing speed (**C**) across sex and time points. Measurements are presented as circles (males) and triangles (females). Mann-Whitney *U* test. Mean ± SEM. *N* = 11 for each genotype and sex. Raw data is available in Supplemental Tables S3 and S4.

**Supplemental Figure S5.**
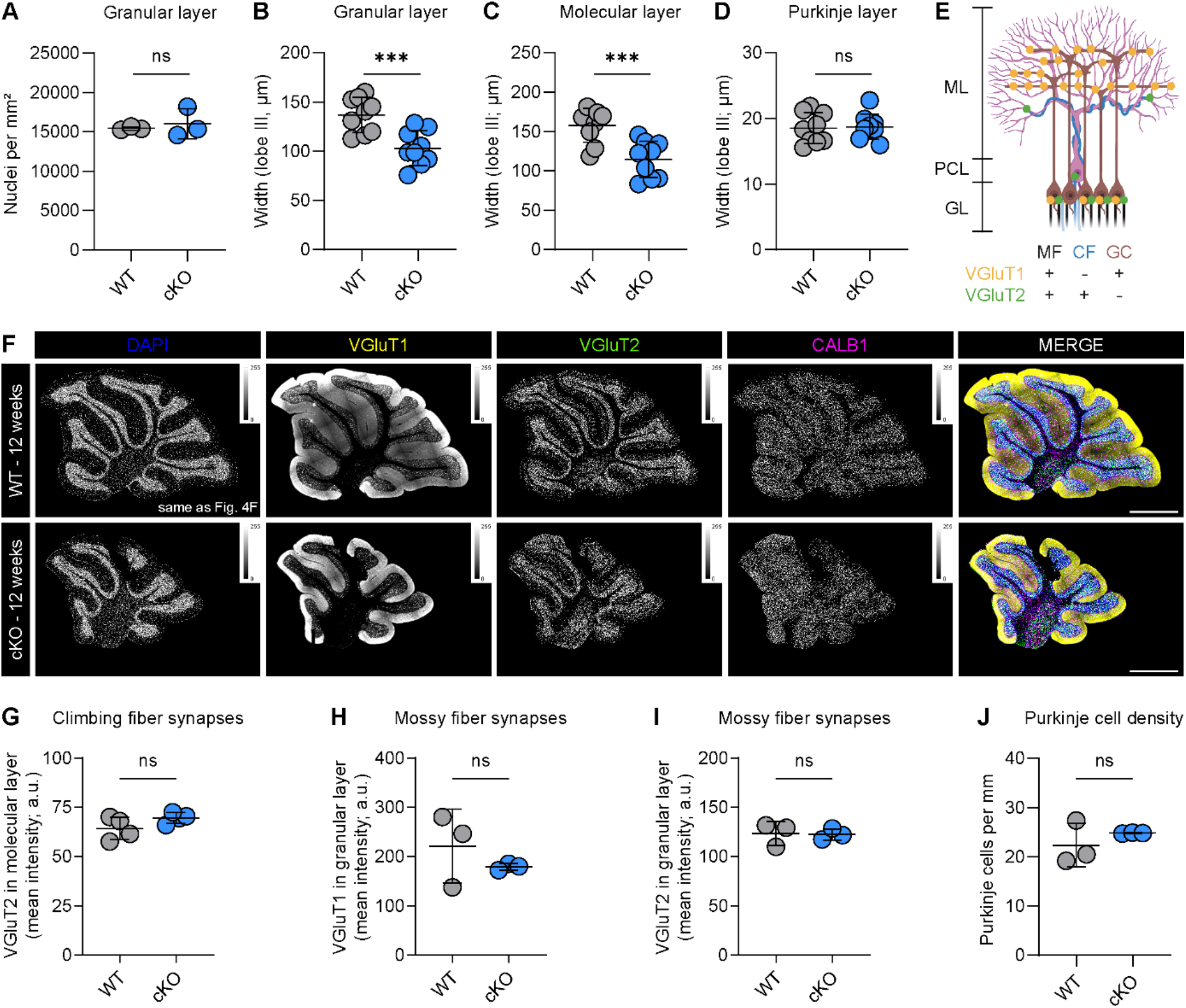
Cerebella of adult *Elp1*^cKO^ mice are smaller. (**A**) Average nuclei number per mm^2^ in granular layer. Granular layer of whole cerebellum was quantified. Unpaired *t*-test. Mean ± SD. *N* = 3. (**B-D**) Average width of granular layer (**B**), molecular layer (**C**) and Purkinje cell layer (**D**) in lobe III. Unpaired *t*-tests. Mean ± SD. *N* = 3 mice/genotype, *n* = 3 sections/mouse with 5 width measurements. (**E**) Schematic illustration of cerebellar microcircuit with Purkinje cell (PC, magenta), granule cells (GC, brown), mossy fiber (MF, black) and climbing fiber (CF, blue) in adult mice. VGluT1 (yellow) is expressed by GCs in molecular layer (ML) and by MFs in granular layer (GL). VGluT2 (green) is expressed by MFs in GL and by CFs in ML and Purkinje cell layer (PCL). Adapted from van der Heijden et al. (2021). (**F**) Representative overview immunofluorescence staining of DAPI (blue), VGluT1 (yellow), VGluT2 (green), CALB1 (magenta) of cerebella of 12-week-old *Elp1*^WT^ and *Elp1*^cKO^ mice. Scale bar, 1 mm. (**G-I**) Mean fluorescence intensity of VGluT2 expression in molecular layer (**G**), VGluT1 (**H**) and VGluT2 (**I**) in granular layer of whole cerebellum overview. Unpaired *t*-tests. Mean ± SD. *N* = ≥ 3 mice/genotype. a.u., arbitrary units. (**J**) Number of Purkinje cells per mm in Purkinje cell layer in whole cerebellum overview. Unpaired *t*-test. Mean ± SD. *N* = 3 mice /genotype.

**Supplemental Figure S6.**
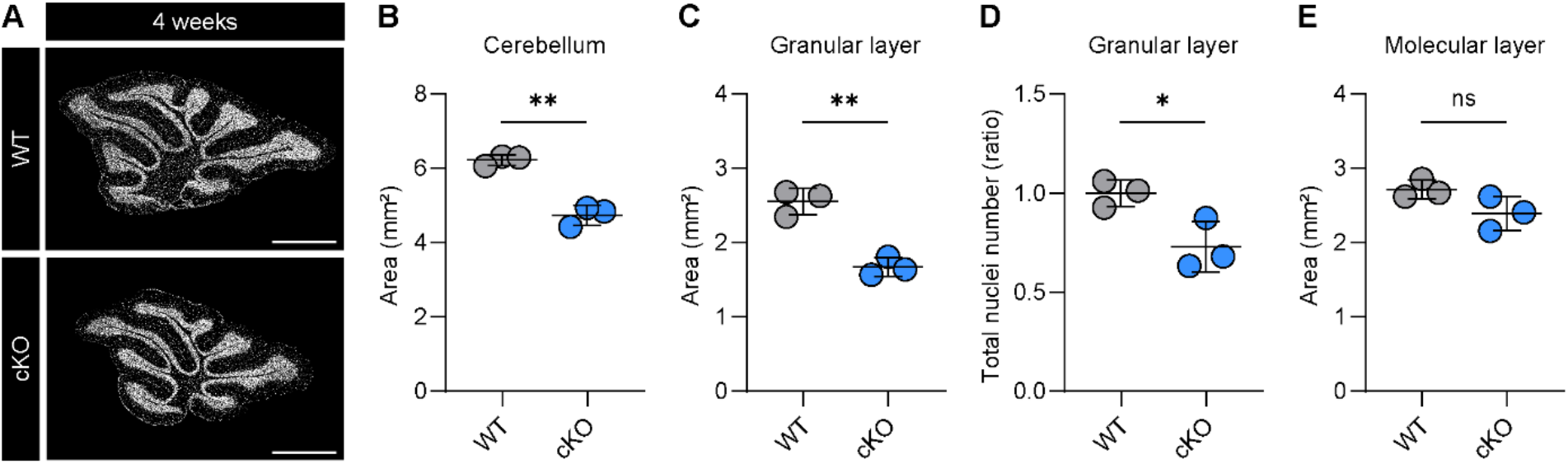
Cerebella of P28 *Elp1*^cKO^ mice are smaller and comprise fewer granule cells. (**A-E**) Representative DAPI staining (**A**) and quantifications (**B-E**) of sagittal sections of *Elp1*^WT^ and *Elp1*^cKO^ whole cerebella at P28. Area of cerebellum (**B**), granular layer (**C**), total nuclei number in granular layer (**D**) and area of molecular layer (**E**) in whole cerebellum overview. Unpaired *t*-tests. Mean ± SD. *N* = 3.

**Supplemental Figure S7.**
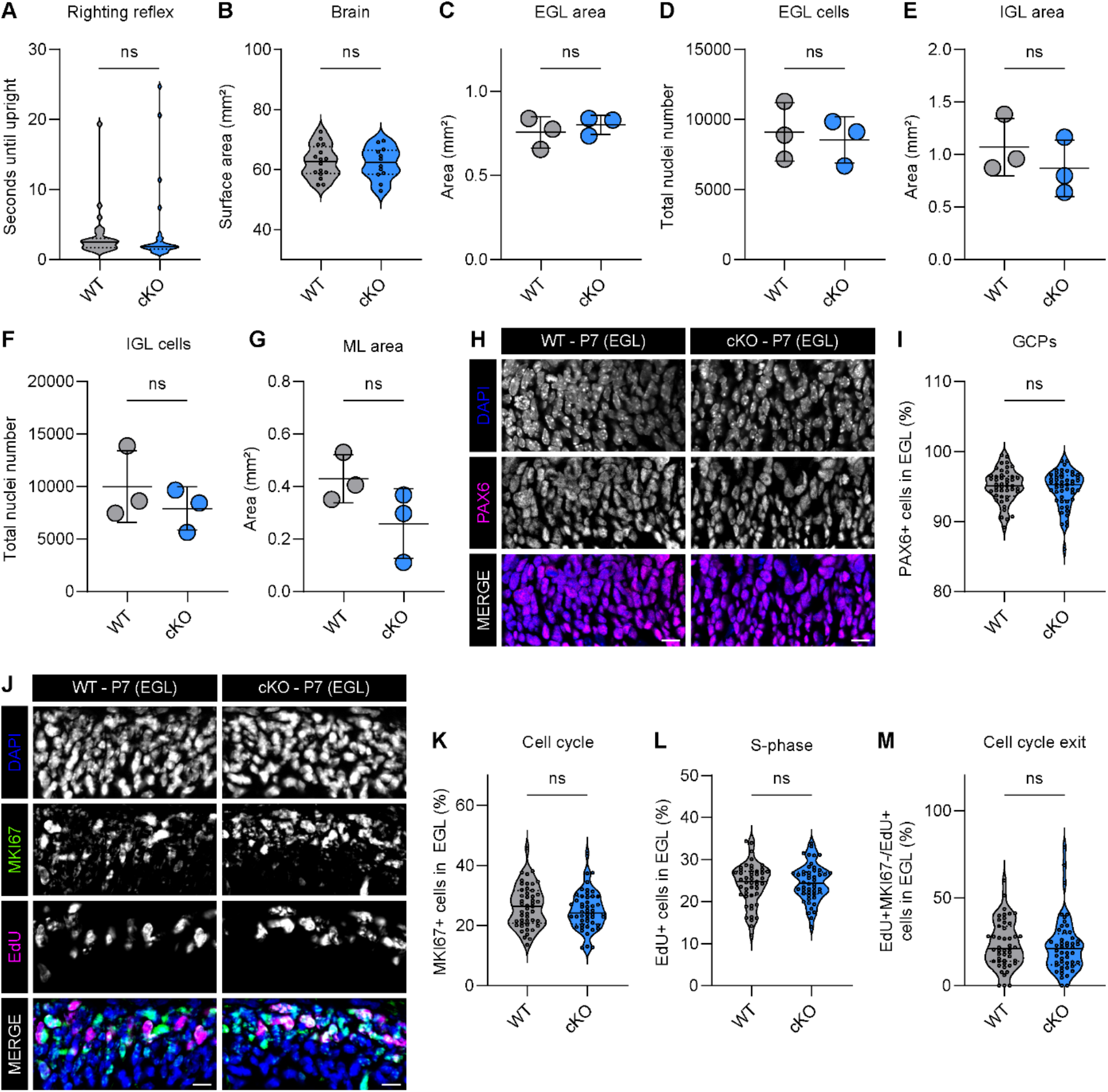
P7 *Elp1*^cKO^ cerebella are smaller but GCPs show similar proliferation rates compared to *Elp1*^WT^ controls. (**A**) Righting reflex of P7 pups. Mann-Whitney *U* test. *N* = 32 (WT) and 31 (cKO). (**B**) Surface area of whole brain from brightfield overview images. Unpaired *t-*tests. *N* = 16 (WT) and 12 (cKO). (**C-G**) Quantifications of sagittal DAPI stainings: area of external granular layer (EGL; **C**), total number of nuclei in EGL (**D**), area of internal granular layer (IGL; **E**), total number of nuclei in IGL (**F**) and area of molecular layer (ML; **G**). Unpaired *t*-tests. Mean ± SD. *N* = 3 mice/genotype. (**H,I**) Representative immunofluorescence staining of GCPs in EGL of lobe VI against PAX6 (**H**) and quantification of PAX6+ cells in EGL of lobes III, VI, IX and X (**I**). Scale bar, 10 μm. Mann-Whitney *U* test. *N* = 3 mice/genotype, *n* = 4 sections/lobe. (**J-M**) Representative immunofluorescence staining of GCPs in EGL against proliferation marker MKI67 and S-phase marker EdU (**J**) and quantifications of MKI67+ (**K**), EdU+ (**L**) and (EdU+MKI67-)/EdU+ cells (**M**) in EGL of lobes III, VI, IX and X. Scale bar, 10 μm. Unpaired *t*-tests. *N* = 3 mice/genotype, *n* = 4 sections/lobe.

**Supplemental Figure S8.**
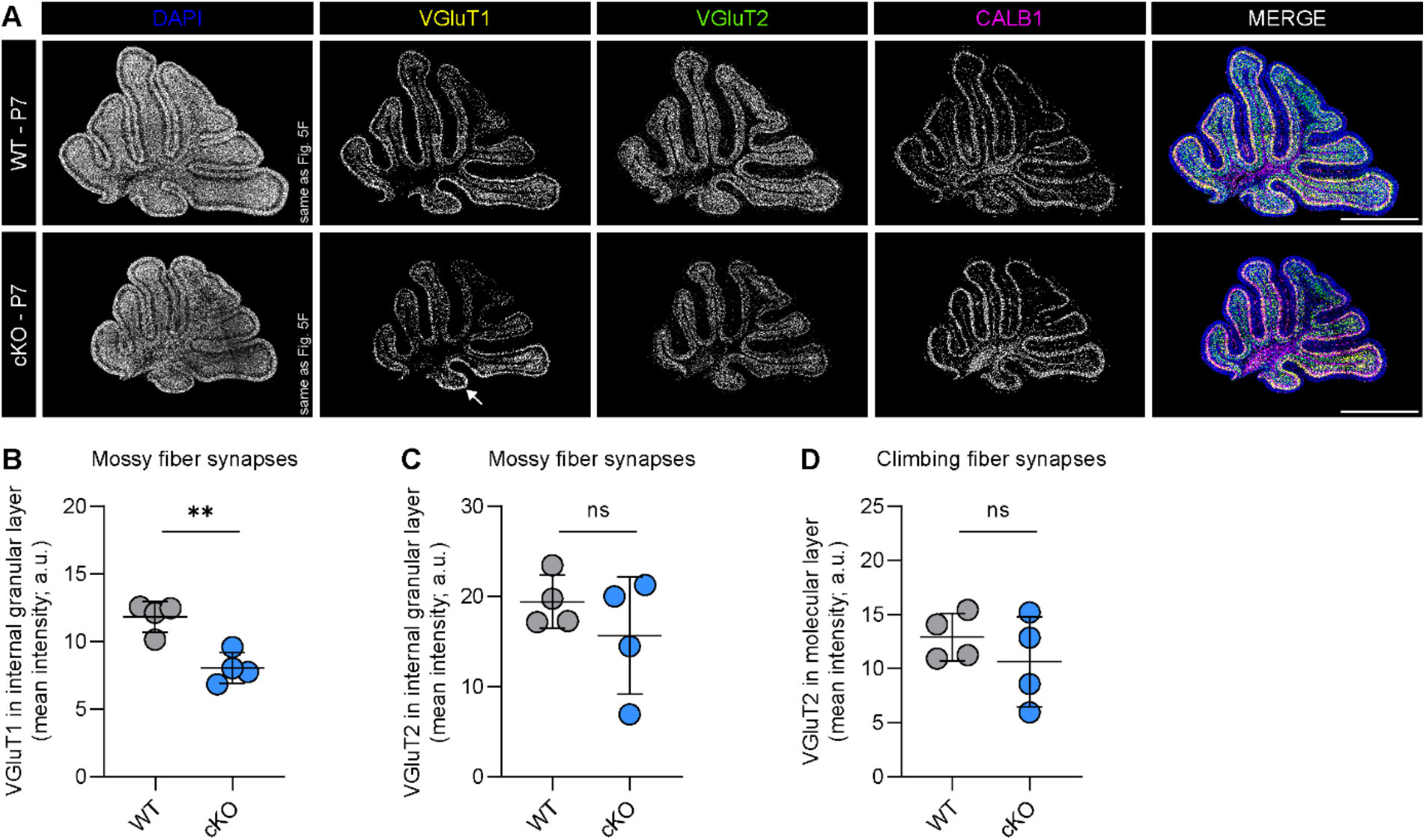
P7 *Elp1*^cKO^ animals express less VGluT1 in molecular and internal granular layer. (**A**) Representative immunofluorescence staining of P7 *Elp1*^WT^ and *Elp1*^cKO^ cerebella against DAPI (blue), VGluT1 (yellow), VGluT2 (green) and CALB1 (magenta). Arrow indicates residual VGluT1 expression in lobe X. Scale bar, 1 mm. (**B-D**) Mean fluorescence intensity of VGluT1 in internal granular layer (mossy fiber synapses; **B**), VGluT2 in internal granular layer (mossy fiber synapses) and VGluT2 in molecular layer (climbing fiber synapses; **D**) in whole cerebellum overview. Unpaired *t*-tests. Mean ± SD. *N* = 4 mice/genotype. a. u., arbitrary units.

**Supplemental Figure S9.**
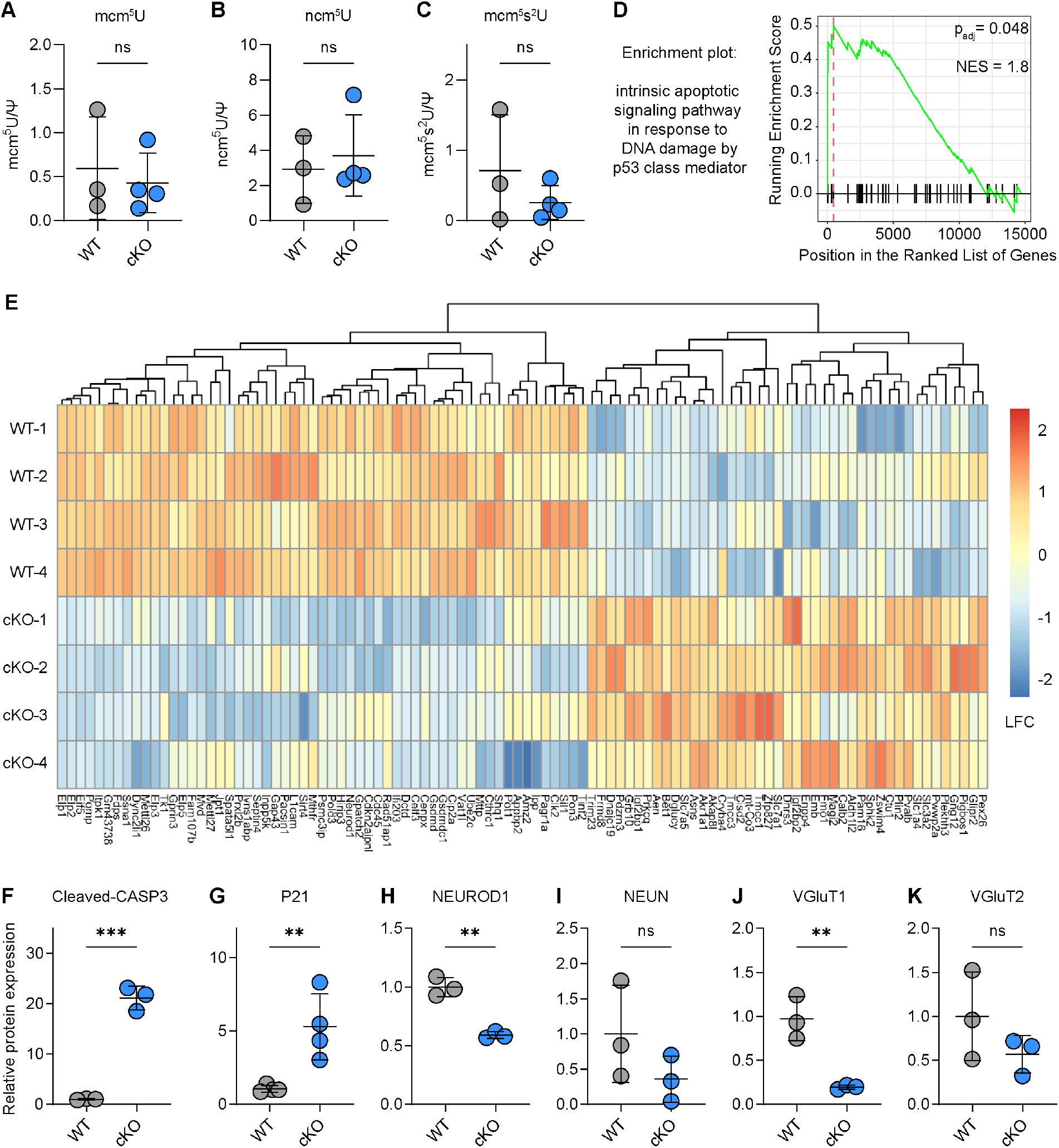
P7 *Elp1*^cKO^ GCPs express higher levels of cell death proteins and lower levels of GC differentiation proteins. (**A-C**) Abundance of Elongator-dependent tRNA modification in P7 *Elp1*^WT^ and *Elp1*^cKO^ GCPs. Modifications were normalized to pseudouridine (Ψ) count. Unpaired *t*-tests. Mean ± SD. *N* ≥ 3 samples/genotype. (**D**) Enrichment plot of gene ontology biological process gene set “intrinsic signaling pathway in response to DNA damage by p53 class mediator” in P7 *Elp1*^cKO^ GCPs. Profile of running enrichment score and positions of gene set members on the rank ordered list are displayed. NES, normalized enrichment score. (**E**) Heat map showing up- (red) and downregulated (blue) proteins in P7 *Elp1*^cKO^ vs *Elp1*^WT^ GCPs. LFC, log fold change. (**F-K**) Immunoblot quantification of protein abundance in GCP lysates normalized to ACTB protein level: cell death marker cleaved-CASP3 (**F**), cell cycle inhibition marker P21 (**G**), GC differentiation markers NEUROD1 (**H**), NEUN (**I**), VGluT1 (**J**) and VGluT2 (**K**). Unpaired *t*-tests. Mean ± SD. *N* = ≥ 3 samples/genotype.

## Supplemental Tables

**Supplemental Table S1**. Primers and antibodies used in the study.

**Supplemental Table S2**. Raw data for behavior analysis, including weight, SHIRPA score, rotarod, and grip strength.

**Supplemental Table S3**. Raw data for catwalk gait analysis of female animals.

**Supplemental Table S4**. Raw data for catwalk gait analysis of male animals.

**Supplemental Table S5**. Significantly up- and down-regulated genes and proteins in *Elp1*^WT^ vs *Elp1*^cKO^ P7 GCPs.

